# CCDC66 regulates primary cilium length and signaling competence via multi-site interactions with transition zone and axonemal proteins

**DOI:** 10.1101/2022.04.10.487777

**Authors:** Deniz Conkar, Ezgi Odabasi, Jovana Deretic, Umut Batman, Kari-Anne M. Frikstad, Sebastian Patzke, Elif Nur Firat-Karalar

## Abstract

The primary cilium is a conserved microtubule-based organelle that serves as a hub for many signaling pathways. It functions as part of the centrosome/cilium complex, which also contains the basal body and the centriolar satellites. Little is known about the mechanisms by which the microtubule-based axoneme of the cilium is assembled with proper length and structure, particularly in terms of the activity of microtubule-associated proteins (MAPs) and the crosstalk between the different compartments of the centrosome/cilium complex. Here, we analyzed CCDC66, a MAP implicated in cilium biogenesis and ciliopathies affecting eye and brain. Live-cell imaging revealed that CCDC66 compartmentalizes between centrosomes, centriolar satellites, and the ciliary axoneme and tip during cilium assembly and disassembly. CCDC66 loss-of-function in human cells causes defects in cilium assembly, length and morphology. Notably, CCDC66 interacts with the MAPs and ciliopathy proteins CEP104 and CSPP1 and cooperates with them during axonemal length regulation. Moreover, CCDC66 interacts with the transition zone protein CEP290 selectively at the centriolar satellites. Its loss disrupts basal body recruitment of transition zone proteins and IFT-B machinery and causes defective Hedgehog signaling. Overall, our results establish CCDC66 as a multifaceted regulator of the primary cilium, and propose a mechanistic insight into how the cooperation of ciliary MAPs as well as subcompartments ensures assembly of a functional cilium.

## Introduction

The primary cilium transduces signaling pathways essential for tissue development and organ homeostasis, including Hedgehog, Wnt and PDGFR signaling (Wheway et al. 2018; Nachury and Mick 2019). Sensory functions of the cilium require its compartmentalization into structural and functional domains as well as crosstalk with the centrosome and centriolar satellites. The primary cilium has a conserved architecture composed of the microtubule-based axoneme and the ciliary membrane, in addition to distinct subcompartments, such as the transition zone, that have variable structures and functions across different organisms (Blacque and Sanders 2014; Lee and Chung 2015). Proximity proteomic studies have identified over 200 proteins as part of the ciliary proteome, although the subciliary localization of the majority is unknown (Mick et al. 2015; Kohli et al. 2017; May et al. 2021). Primary cilium biogenesis is a tightly-regulated, multistep process that involves centriolar maturation to a basal body, elongation of axonemal microtubules from the basal body, formation of the ciliary membrane, and establishment of the transition zone as the ciliary gate (Mirvis et al. 2018; Breslow and Holland 2019). Precise spatiotemporal control of its assembly and disassembly kinetics, length, stability, structure and composition is required to ensure proper cilium function. As such, deregulation of these processes causes various human diseases including the multisystem pathologies of the eye, kidney, skeleton, brain and other organs, collectively named “ciliopathies” (Braun and Hildebrandt 2017; Reiter and Leroux 2017). Defining the complex mechanisms by which a functional cilium is built and maintained is essential for providing insight into the molecular defects that underlie ciliopathies and their phenotypic heterogeneity.

The transition zone functions as the ciliary gate to control selective ciliary entry and exit of cargoes and forms structural links between the axoneme and the ciliary membrane (Szymanska and Johnson 2012; Goncalves and Pelletier 2017). Mutations affecting transition zone proteins are prevalent in ciliopathies, highlighting the importance of understanding its biogenesis and function. The transition zone cooperates with multiple protein complexes and cellular structures to regulate cargo trafficking. Intraflagellar transport (IFT)-A and IFT-B machineries and the BBSome complex traffic ciliary proteins along the axoneme between the ciliary base and the tip. (Nachury et al. 2010; Pedersen et al. 2012; Garcia-Gonzalo and Reiter 2017; Nachury and Mick 2019). Additionally, membrane-less granules that move around the primary cilium, known as centriolar satellites, control cilium composition by regulating ciliary protein targeting (Odabasi et al. 2020; Prosser and Pelletier 2020). They have been proposed to act as trafficking machines for centrosome/cilium proteins such as IFT-B components (Odabasi et al. 2019; Aydin et al. 2020). Another emerging mechanism for cilium content regulation is ectocytosis of ciliary proteins from the cilium tip, which is the region between the ciliary membrane and the plus ends of the furthest reaching ciliary microtubules (Wang and Barr 2016; Nager et al. 2017; Phua et al. 2017). The cilium tip has also been implicated in Hedgehog signaling, IFT turnover and remodeling of ciliary microtubules (He et al. 2014; Pedersen and Akhmanova 2014; Chien et al. 2017; Conkar and Firat-Karalar 2020). Despite its critical functions, the tip region remains one of the most poorly characterized ciliary subcompartments with respect to its biogenesis and composition.

The ciliary axoneme provides structural support to the cilium and serves as the track for bidirectional transport of cargoes. It is composed of nine radially-arranged, remarkably stable doublet microtubules, which are highly modified by posttranslational modifications (PTMs) (Wloga et al. 2017). The plus ends of axonemal microtubules are spatiotemporally regulated in order to maintain proper cilium structure and length. However, relatively little is known about the mechanisms by which axonemal microtubules of the primary cilium are nucleated and elongated from the centriolar template at the right length. Given the critical functions of microtubule-associated proteins (MAPs) in spatiotemporal regulation of microtubule dynamics, identification and characterization of ciliary MAPs is required to address these questions.

Several microtubule-associated proteins (MAPs) mutated in ciliopathies have been characterized as components of the ciliary tip and the axoneme, which provide leads to dissect axonemal assembly and organization (Bodakuntla et al. 2019; Conkar and Firat-Karalar 2020). For example, CEP104 interacts with CSPP1 and functions during assembly of Hedgehog-competent cilia at the right length (Frikstad et al. 2019). Notably, CEP104 contains a tubulin-binding TOG domain, which promotes microtubule polymerization *in vitro* and is required during ciliary length regulation (Satish Tammana et al. 2013; Das et al. 2015; Rezabkova et al. 2016; Al-Jassar et al. 2017; Yamazoe et al. 2020). Finally, CEP104, together with CCDC66, ARMC9, TOGARAM1 and CSPP1, is part of a protein module mutated in the ciliopathy Joubert syndrome, suggesting that they might work together during cilium biogenesis and function (Latour et al. 2020). Timing and dynamics of their localization to the primary cilium, whether they form functional complexes at cilium, and the nature of their relationship with the axonemal microtubules remain poorly understood.

We previously identified CCDC66 as a MAP that localizes to the centrosome, centriolar satellites and primary cilium, and regulates cilium formation (Sharp et al. 2011; Gupta et al. 2015; Conkar et al. 2017). CCDC66 also stably localizes to the ciliary axoneme, suggesting that it might be a structural component (Conkar et al. 2019). Early frameshift mutations of CCDC66 in dogs and its deletion in mouse cause retinal degeneration and olfactory deficits (Dekomien et al. 2010; Gerding et al. 2011; Schreiber et al. 2018; Murgiano et al. 2020), and it was also identified as part of a Joubert module, although no Joubert-causative *CCDC66* mutations are hitherto not reported (Latour et al. 2020) Together, these findings suggest that CCDC66 is an important regulator of primary cilium structure and/or function. However, its precise ciliary functions and molecular mechanisms of action are unknown.

Here, we used localization, interaction studies and loss-of-function experiments to define the ciliary functions and mechanisms of CCDC66. High resolution and live imaging experiments identified CCDC66 as a new component of the ciliary axoneme/tip and revealed its dynamic compartmentalization at the cilium, basal body and centriolar satellites during cilium assembly and disassembly. Furthermore, CCDC66 interacts with the transition zone protein CEP290 at the centriolar satellites and the axonemal proteins CEP104 and CSPP1 at the cilia. By ensuring proper ciliary recruitment of these proteins, CCDC66 regulates cilium assembly, length and signaling. Our results identify CCDC66 as a new ciliary MAP required for cilium structure and function, and advance our understanding of the coordinated activity of CCDC66 with other MAPs at the axoneme and centrosomal/ciliary subcompartments.

## Results

### CCDC66 localizes to the ciliary axoneme and tip and exhibits highly dynamic localization behavior during ciliogenesis

To examine the localization of CCDC66 relative to centrosomal and ciliary subcompartments, we induced ciliogenesis in the previously characterized retinal pigment epithelial cells stably expressing GFP-CCDC66 (from now on: RPE1::GFP-CCDC66) by serum starving them for 24 h, and stained with antibodies against proteins at the distal centriole (Centrin), distal appendages (CEP164), transition zone (CEP290), IFT-B machinery (IFT88), ciliary membrane (ARL13B) and axoneme (acetylated tubulin) (Conkar et al. 2017). Plot profile analysis was performed along the axis of the centrosome and cilium and comparing the signal intensity of CCDC66 relative to different markers (Fig. 1A). CCDC66 localized proximal to CEP164 and Centrin at the basal body. CCDC66 did not co-localize with CEP290 at the transition zone, instead they co-localized at the granules around the centrosome. At the primary cilium, CCDC66 localized to the axoneme in a punctate manner, in contrast to the relatively homogenous staining of the ciliary membrane by the small GTPase ARL13B (Fig. 1A). Its relative localization to IFT88 and acetylated tubulin revealed enrichment of CCDC66 at the ciliary tip in a fraction of cells, which identifies CCDC66 as a new tip protein (Fig. 1A). To determine the localization for endogenous CCDC66, we tried two commercially available antibodies raised against human CCDC66 and one custom antibody raised against mouse CCDC66. However, none of the antibodies detected endogenous CCDC66 at the cilia in human cell lines.

**Figure 1.**
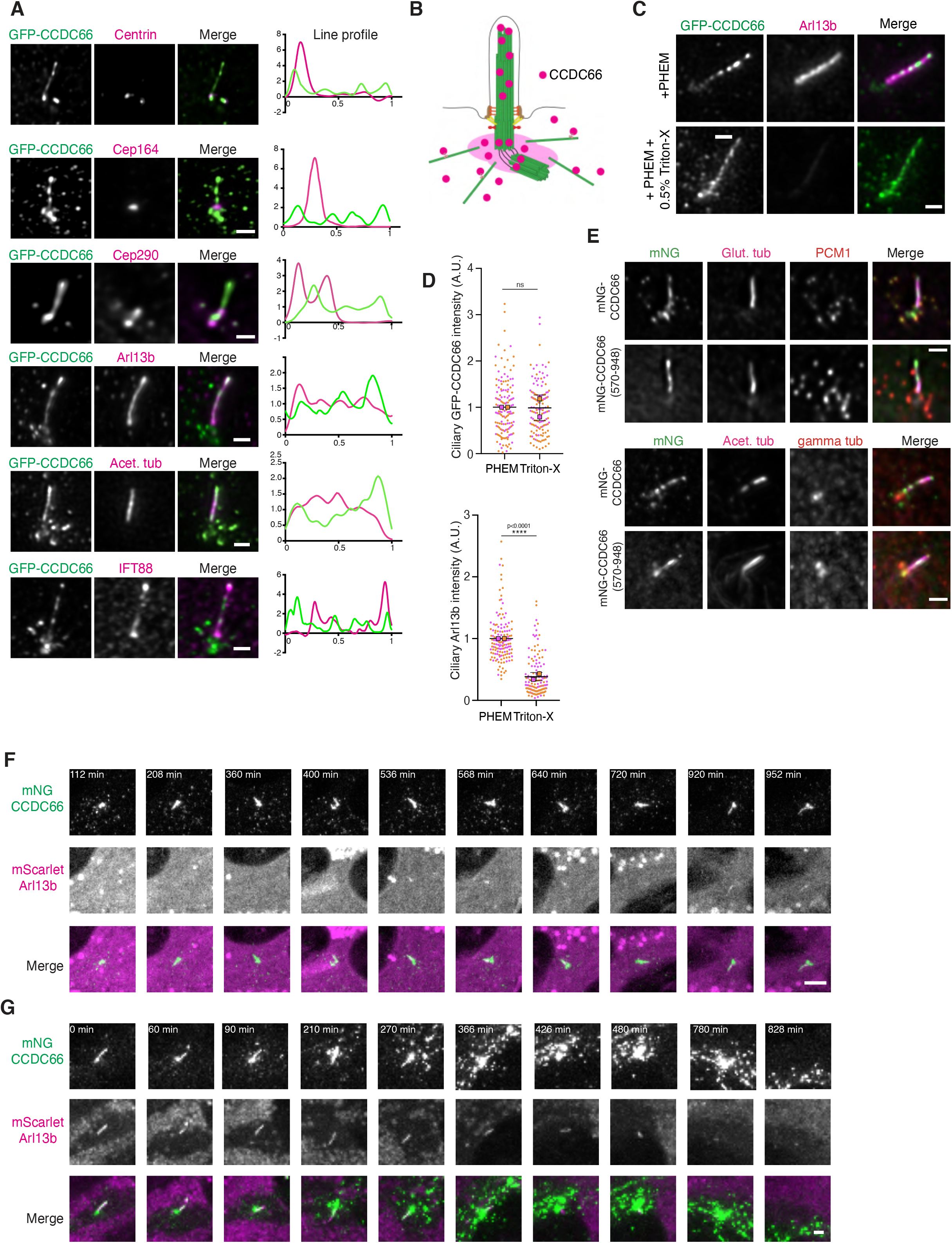
CCDC66 exhibits highly dynamic spatiotemporal dynamics during cilium assembly and disassembly. **(A) Sub-centrosomal and -ciliary localization of CCDC66 in ciliated cells**. RPE1::GFP-CCDC66 cells were serum starved for 48 hours, fixed and stained for GFP along with centriole (centrin), distal appendage (CEP164), transition zone (CEP290) and primary cilium (ARL13B, acetylated tubulin and IFT88) markers. Scale bar: 5 μm for acetylated tubulin co-localization, 1 μm for the rest of the images. **(B) Schematic representation of CCDC66 localization at the centrosome/cilium complex**. CCDC66 localizes to the ciliary axoneme/tip, the basal body and the centriolar satellites. It is excluded from the transition zone at the primary cilium. **(C-D) CCDC66 stably associates with the ciliary axoneme. (C)** Ciliated RPE1::GFP-CCDC66 cells were treated with PHEM or PHEM + 0.5% Triton-X for 30 seconds, fixed with PFA and stained for GFP and ARL13B. **(D)** Ciliary GFP-CCDC66 and ARL13B intensities were quantified by measuring corresponding ciliary intensities and subtracting the background signal. Magenta and orange represent individual values from two independent experiments. Scale bar: 5μm (70 cilia/experiment, ****P < 0.0001, ns: not significant t-test) **(E) C-terminal microtubule binding fragment (570-948 a.a.) of CCDC66 localizes to the basal body and the axoneme**. RPE1::mNeonGreen (mNG)-CCDC66 and RPE1::mNG-CCDC66 (570-948) cells were serum starved for 48 hours, fixed and stained for mNeonGreen, glutamylated tubulin and PCM1 or acetylated tubulin and gamma tubulin antibodies. Scale bar: 2μm. **(F) Spatiotemporal dynamics of CCDC66 during cilium assembly**. RPE1::mNG-CCDC66, mScarlet-ARL13B cells were imaged with confocal microscopy at 8 minute intervals after serum withdrawal. First time point (112 min) indicates the starting of cilium formation. Scale bar: 5μm **(G) Spatiotemporal dynamics of CCDC66 during cilium disassembly**. RPE1::mNG-CCDC66, mScarlet-ARL13B cells were serum starved for 48 hours and imaged with confocal microscopy at 6 minute intervals after serum addition. Scale bar: 2μm

Taken together, CCDC66 has co-existing cellular pools, at the basal body, the primary cilium (axoneme and ciliary tip), and at the centriolar satellites, suggesting multiple functions at the primary cilium (Fig. 1B)

CCDC66 directly binds to microtubules, localizes to the ciliary axoneme and its ciliary pool is immobile (Conkar et al. 2017; Conkar et al. 2019). These findings led us to hypothesize that it might be a structural component of the axoneme To test this, we examined CCDC66 localization in serum starved RPE1::GFP-CCDC66 cells treated with 0.5% Triton-X, which removes the surrounding ciliary membrane and membrane-associated proteins including ARL13B from the axoneme (Nachury et al. 2007). As compared to control cells, detergent treated cells had about 3-fold reduction in their ciliary ARL13B intensity (p<0.0001), while ciliary GFP-CCDC66 intensity remained relatively unchanged (Fig. 1C, 1D). Given its axonemal association, we next asked whether the microtubule-binding activity of CCDC66 is required the ciliary localization of CCDC66. In prior work, we showed that C-terminal 570-948 residues of CCDC66 binds to MTs *in vitro* (Conkar et al. 2017). To test whether this fragment targets CCDC66 to the cilia, we generated RPE1 cell lines stably expressing mNeonGreen (mNG) fusions of CCDC66 and CCDC66 (570-948). Both mNG-CCDC66 and mNG-CCDC66 (570-948) localized to the basal body, axoneme and the ciliary tip (Fig. 1E). In contrast to mNG-CCDC66, mNG-CCDC66 (570-948) did not localize to centriolar satellites (Fig. 1E). Thus, CCDC66 is stably associated with the axoneme in the primary cilium and its microtubule-binding fragment is sufficient for targeting it specifically to the basal body and the primary cilium.

To determine how CCDC66 functions in ciliogenesis, we monitored its spatiotemporal localization dynamics in RPE1::mNG-CCDC66 cells transduced with the ciliary membrane marker mScarlet-ARL13B and induced for cilium assembly by serum starvation (Fig.1F and Movie 1). We chose mNG over GFP as it is brighter and more photostable (Shaner et al. 2013). In unciliated cells, mNG-CCDC66 localized to the centriolar satellites and centrosome. Following serum starvation, mNG-CCDC66 localized to the developing cilia about the same time as mScarlet-ARL13B. As they were recruited to the growing ciliary axoneme, the number of CCDC66-positive centriolar satellite granules gradually decreased and became less concentrated around the basal body (Fig.1F). All ciliated cells exhibited ciliary localization of CCDC66 after ARL13B-positive cilia formed, and CCDC66 was retained at the cilia after its recruitment. Analogous to its localization profile in fixed cells, ciliary mNG-CCDC66 was heterogeneously distributed along the cilia and enriched at the cilium tip. Growing or steady state GFP-CCDC66-postive cilia were present in about 80% cells after 24 h serum starvation. Notably, the timing of initiation of ciliogenesis as well as the duration between initiation and formation of steady state cilia after serum starvation varied from cell to cell (Fig. S1A). Ciliary recruitment of CCDC66 at the early stages of cilium formation and reduction of its satellite pool concomitant to the initiation and growth of the cilium is further supported by immunofluorescence analysis of RPE1::mNG-CCDC66 cells fixed at different time points after serum starvation and stained for the axoneme, ciliary membrane and satellites (Fig. S1B, S1C). Thus, CCDC66 localizes to the primary cilium during the initial stages of cilium formation and is a constitutive resident of the cilium and the cilium tip.

We next monitored CCDC66 localization dynamics during cilium disassembly. RPE1::mNG-CCDC66 / mScarlet-ARL13B cells were serum starved for 48 hours and imaged following serum addition (Fig. 1G and Movie 2). Consistent with described mechanisms of cilium disassembly (Mirvis et al. 2019), the ciliary pools of CCDC66 and ARL13B were lost by multiple different events including ciliary decapitation, resorption of the axoneme and whole-cilium shedding (Fig. 1G, S1D-E). Notably, in the reverse order as that seen during cilium assembly, as the axonemal pool of CCDC66 disappeared, the centriolar satellite pool reappeared, suggesting possible redistribution of CCDC66 from cilium to the satellites. We also observed that CCDC66 accumulated at the centrosome during cilium disassembly, supporting its relocation. This is unlikely due to GFP-CCDC66 expression level in this cell line, since we previously showed that it was expressed at physiological levels (Conkar et al. 2017). Immunofluorescence analysis of cells fixed and stained at different time points after serum stimulation confirmed these observations (Fig. S1E). Collectively, our results show that CCDC66 enters the primary cilium early during its assembly, stably localizes to the axoneme and ciliary tip, and exits the cilium during its disassembly.

### CCDC66 is required for assembling primary cilium with proper length

Our previous work indicated a role of CCDC66 during primary cilium assembly (Conkar et al. 2017). However, the precise roles of CCDC66 during cilium assembly and maintenance is unknown. To address this, we first investigated how CCDC66 affects the kinetics of cilium biogenesis using siRNA-mediated loss-of-function experiments. We quantified the percentage of control and CCDC66-depleted cells that formed cilia over a 48 h serum starvation time course by staining with acetylated tubulin (Fig. 2A, Fig. S2A-B). The fraction of ciliated cells was reduced upon CCDC66 depletion at all time points after serum starvation and did not reach control levels even at 48 h post serum starvation (Fig. 2B). Additionally, we stained cells with proliferation marker Ki-67 and found that the percentage of quiescent cells (Ki-67 negative cells) between control and CCDC66-depleted RPE1 cells was similar (Fig. S2C, 2D). This data shows that failure to enter quiescence does not account for the defective ciliation of CCDC66-depleted cells. Finally, we quantified the effects of CCDC66 loss on cilium length. Cilia that formed in CCDC66-depleted cells were significantly shorter relative to control cells at all time points following serum starvation (Fig. 2C). Thus, CCDC66 is required for both the initiation of cilium assembly and elongation of the axoneme.

**Figure 2.**
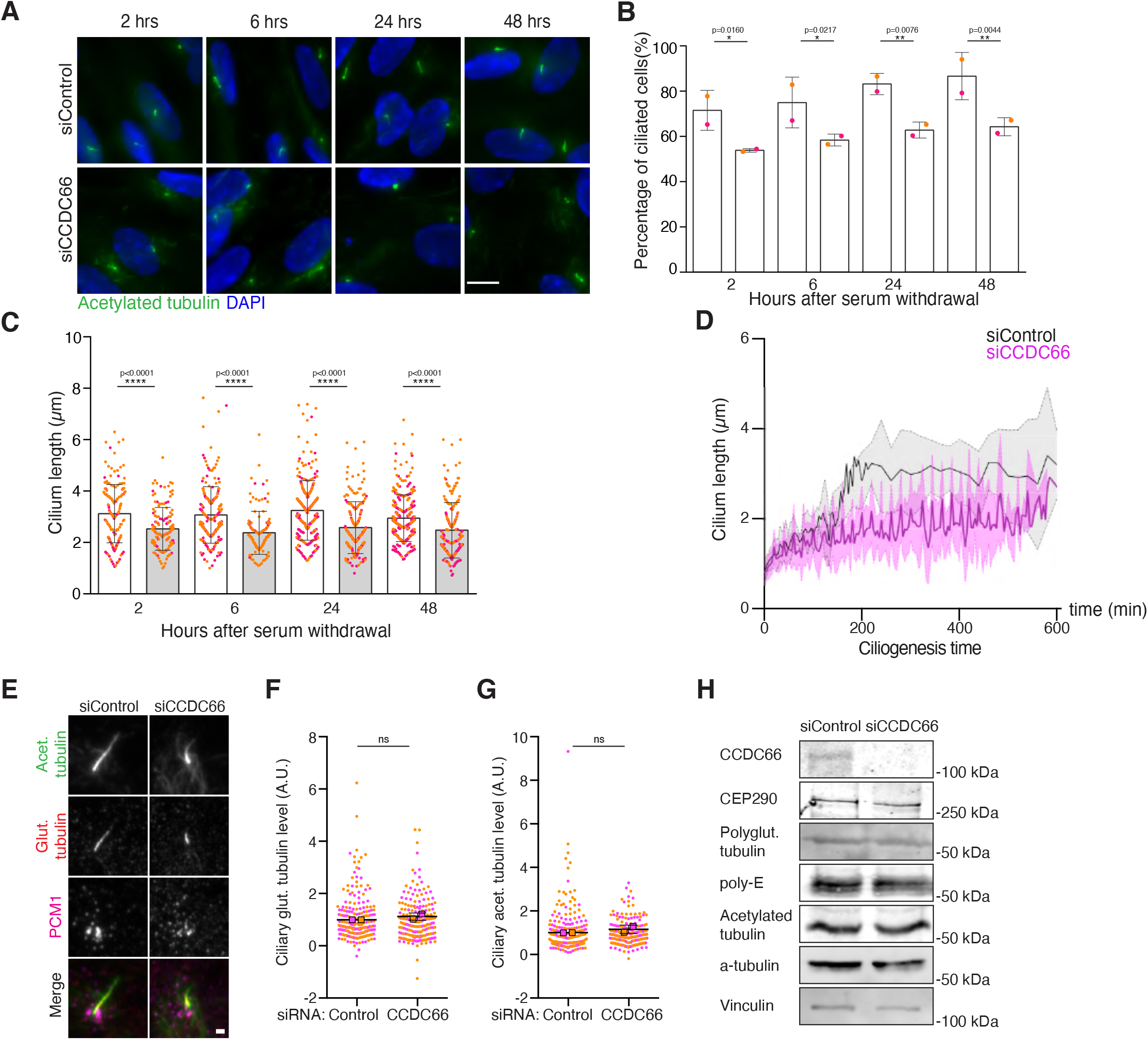
CCDC66 is required for efficient ciliogenesis and axoneme elongation. **(A-B) CCDC66 loss impairs cilium formation. (A)** Control and CCDC66 depleted RPE1 cells were fixed at 2, 6, 24 and 48 hours after serum starvation. Following fixation, cells were stained for anti-acetylated tubulin antibody and DAPI. **(B)** Percentage of cilium formation was quantified by dividing the cilium number determined by counting acetylated tubulin by total cell number determined by counting nuclei and plotted against time after serum starvation. Magenta and orange represent individual values from two independent experiments. Error bars represent ±SD. (100 cells/serum starvation time point for each experiment, *P < 0.05, **P < 0.01 two way ANOVA) Scale bar: 10μm **(C) CCDC66 depletion leads to shorter cilia**. Ciliary length from (A) was measured and plotted against time after serum starvation. Magenta and orange represent individual values from two independent experiments. Error bars represent ±SD. (100 cilia/serum starvation time point for each experiment, *P < 0.05, **P < 0.01 two way ANOVA) **(D) CCDC66 depletion results in defective axoneme elongation**. RPE1::mCitrine-SMO cells were transfected with two rounds of control or CCDC66 siRNAs. 48 h post-transfection, they were monitored by time lapse imaging over 24 h every 4 min. Lengths of cilia were measured from two independent experiments as SMO-positive cilia appeared (t=0). Data represent the mean ±SD from 119 (siControl) and 101 (siCCDC66) cilia of two independent experiments. **(E-G) CCDC66 loss does not alter ciliary acetylation and polyglutamylation. (E)** RPE1 cells were transfected with two rounds of control or CCDC66 siRNA and serum starved for 48 hours. Cells were fixed and stained for acetylated tubulin, polyglutamylated tubulin and PCM1 antibodies. **(F)** Ciliary acetylated tubulin and **(G)** polyglutamylated tubulin levels were quantified from **(E)** by measuring the PTM intensities via subtracting the background signal, multiplied by the signal area, and dividing by the cilium length. Data represent the mean ±SD. Magenta and orange represent individual values from two independent experiments. (100 cilia/experiment, ns: not significant, t-test) Scale bar: 1μm **(H) Effects of CCDC66 depletion on cellular abundance of various proteins**. RPE1 cells were transfected with two rounds of control or CCDC66 siRNA and serum starved for 48 hours. Cell lysates were prepared, resolved on SDS-PAGE and immunoblotted for CCDC66, CEP290, polyglutamylated tubulin, poly-E, acetylated tubulin, alpha-tubulin and vinculin (loading control).

To spatiotemporally define the cilium elongation defects, we performed live imaging of control and CCDC66 siRNA-transfected RPE1 cells stably expressing the mCitrine fusion of the ciliary transmembrane protein Smoothened (mCitrine-SMO) during cilium assembly (Fig. 2D). After the cells committed to cilia formation, cilia grew slower in CCDC66-depleted cells relative to control cells. By 200 min, cilium growth plateaued in CCDC66-depleted cells but not in control cells. By 10 h, the average length of cilia in control cells and CCDC66 depleted cells were about 3.2 μm and 2.7 μm, respectively (Fig. 2D). Notably, the kinetics of cilia growth and shortening at steady state was nearly balanced in control cells, while there were high length variations in CCDC66-depleted cells (Fig. 2D). We also examined the behavior of steady state cilia using live imaging by quantifying the following two events from videos: 1) ectocytosis or decapitation from cilium tips, 2) breakage (scission), which represents the thinning of the distal axoneme followed by its breakage, 3) ripping off, where the cilium starts to elongate abnormally and rips off from a point of enriched mCitrine-SMO signal (Fig. S2E). Percentage of ciliary ectocytosis/decapitation decreased from 67% in control ciliated cells to 60% in CCDC66-depleted ones (Fig. S2F). CCDC66-depleted cilia also showed an increase in the frequency of tip breakage and scission. Although these differences might also underlie the shorter cilium phenotype, they were not statistically significant.

Cilium assembly and maintenance require remodeling of the microtubule cytoskeleton by a diverse array of MAPs and tubulin PTMs (Pedersen et al. 2012; Broekhuis et al. 2013; Keeling et al. 2016). To examine whether CCDC66 loss results in defective modification of the ciliary tubulins, we quantified ciliary polyglutamylated and acetylated tubulin levels and found that their levels was similar in control and CCDC66-depleted cells (Fig. 2E-G). Cellular levels of polyglutamylated and acetylated tubulin were comparable between control and CCDC66 depleted cells (Fig. 2H). Taken together, these findings identify CCDC66 as a regulator of cilium length.

### CCDC66 depletion interferes with transition zone recruitment of CEP290 and and basal body recruitment of the IFT-B machinery

Cilium assembly is a multistep process that requires the coordinated activity of a multitude of proteins, cellular structures and signaling pathways (Mirvis et al. 2018; Breslow and Holland 2019). To investigate how CCDC66 regulates ciliogenesis, we used quantitative immunofluorescence to analyze whether known regulators of different stages of ciliogenesis are properly recruited to the centrosomes or cilia upon RNAi-mediated knockdown of CCDC66. To this end, control and CCDC66-depleted cells were serum starved for 48 h and stained for antibodies against components of distal appendages, transition zone proteins and IFT machinery (Fig. 3B-S).

**Figure 3.**
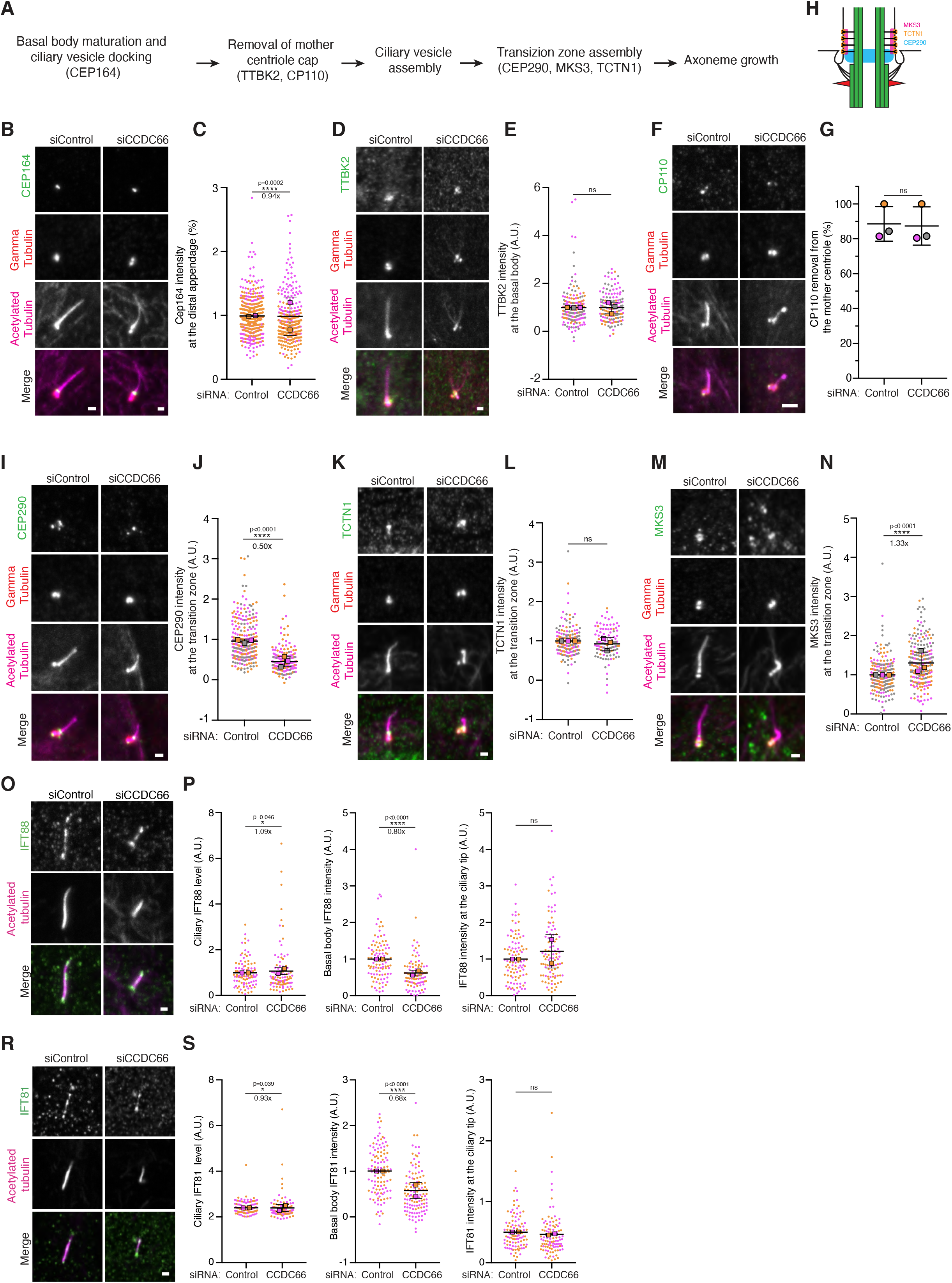
CCDC66 is important for proper transition zone formation and IFT-B localization to the basal body. **(A)** Stepwise assembly of the primary cilium and cartoon representation of 3 transition zone modules. **(B-N) CCDC66 loss impairs CEP290 and MKS3 levels at the transition zone**. RPE1 cells were transfected with two rounds of control or CCDC66 siRNA and serum starved for 48 hours. Cells were fixed and stained for **(B)** CEP164, **(D)** TTBK2, **(F)** CP110, **(I)** CEP290, **(K)** TCTN1, or **(M)** MKS3, and gamma-tubulin and acetylated tubulin antibodies. **(H)** Cartoon representation of 3 transition zone modules. CEP164 levels at the distal appendages, basal body TTBK2 levels, CP110 removal from the mother centriole, CEP290, TCTN1 and MKS3 levels at the transition zone were quantified and plotted. To quantify only transition zone pool of the protein, gamma tubulin is taken as reference. The pools of the CEP290, TCNT1 and MKS3 above the gamma tubulin are quantified as transition zone. Data represent the mean ±SD. Magenta, orange and grey represent individual values from three independent experiments. Scale bar: 2μm for B, D and F, 1μm for I, K and M (100 cells/experiment, ****P < 0.0001, ns: not significant t-test). **(O-S) Basal body levels of IFT-B components decrease upon CCDC66 depletion**. RPE1 cells were transfected with two rounds of control or CCDC66 siRNA and serum starved for 48 hours. Cells were fixed and stained for **(O)** IFT88 and **(R)** IFT81 and acetylated tubulin antibodies. **(P-S)** Ciliary, basal body and ciliary tip levels of IFT88 and IFT81 are plotted. Ciliary IFT81 and IFT88 levels were quantified by measuring the protein intensity at cilia, subtracting the background signal, multiplied by the signal area, and dividing by the cilium length. Ciliary IFT81 and IFT88 levels were normalized relative to the mean of control group.(=1) Data represent the mean ±SD. Magenta and orange represent individual values from two independent experiments. (100 cells/experiment, *P < 0.01, ****P < 0.0001, ns: not significant t-test). Scale bar: 2μm

During ciliogenesis, distal appendage protein CEP164 recruits the kinase TTBK2, which phosphorylates CEP83 and MMP9 and results in removal of the centriole capping protein complex CP110-CEP97 from the mother centriole (Graser et al. 2007; Tanos et al. 2013; Cajanek and Nigg 2014). While CCDC66 depletion resulted in a minor 6 percent reduction centrosomal CEP164 levels, it did not alter centrosomal TTBK2 levels (Fig. 3B - E). There was also no defect in the removal of CP110 from the mother centriole upon CCDC66 loss (Fig. 3F, 3G). These results indicate that CCDC66 is not required for acquisition of distal appendages or removal of the centriole cap.

Centriole cap removal is followed by periciliary vesicles docking to the basal body and their fusion, formation of the transition zone and elongation of the axoneme (Breslow and Holland 2019). We previously showed that CCDC66 interacts with the transition zone protein CEP290, which plays critical roles in transition zone assembly and ciliary content regulation (Betleja and Cole 2010; Drivas and Bennett 2014). The transition zone consists of the membrane-associated or non-membranous NPHP, MKS and CEP290 modules that form hierarchically (Fig. 3H) (Garcia-Gonzalo and Reiter 2017; Goncalves and Pelletier 2017). To assess how CCDC66 loss affects these modules, we quantified the basal body levels of one protein from each module, namely TCTN1, CEP290 and MKS3 (Fig. 3I-N). While CCDC66 depletion did not alter basal body TCTN1 levels (Fig. 3K, 3L), there was about 0.5-fold decrease in CEP290 (Fig. 3I, 3J) and 1.3-fold increase in MKS3 levels at the basal body (Fig. 3M, 3N). Immunoblotting revealed similar cellular levels of CEP290 between control and CCDC66-depleted cells, indicating that CCDC66 regulates CEP290 targeting to the transition zone (Fig. 2H). These defects suggest that CCDC66 might take part in transition zone assembly by governing proper targeting of specific proteins to the transition zone.

After transition zone assembly, the axoneme is templated from the centriolar base and the mature cilium is assembled. This process requires proper activity of the IFT machinery, which traffics proteins such as tubulin dimers along the ciliary axoneme. Therefore, we examined the recruitment of the IFT-B machinery to the basal body and cilium as putative mechanisms that underlie the ciliary defects of CCDC66-depleted cells (Fig 3O-S). To this end, we measured the levels of the IFT-B components IFT88 and IFT81 at the cilium ciliary tip and basal body. While the ciliary tip levels of IFT88 and IFT81 remained unaltered, their basal body levels but not ciliary levels were impaired in CCDC66-depleted cells (Fig 3P, 3S). There results identify a function for CCDC66 during basal body recruitment of the IFT-B machinery.

### Basal body and axonemal pools of CCDC66 are required for its functions during cilium and transition zone assembly

To address whether and how the ciliary functions of CCDC66 are governed by its distinct cellular pools (at the basal body, centrosomes and primary cilium), we performed phenotypic rescue experiments with three different CCDC66 siRNA-resistant mutants (Fig. 4A): 1) mNG-CCDC66 to validate the specificity of the phenotypes, 2) mNG-CCDC66 (570-948) to assess the functional significance of CCDC66 localization to the basal body and axoneme, 3) mNG-CCDC66 fusion fwith the centrosomal localization sequence of AKAP450 (PACT domain) at its C-terminus to distinguish its centrosome-specific activities from the ones mediated by satellites and the axoneme. For these experiments, we generated RPE cell lines stably expressing only mNG as a control along with mNG fusions of CCDC66 mutants (Fig. S3A). Additionally, we examined localization of the fusion proteins in serum-starved, CCDC66-depleted stable cells stained for acetylated tubulin and gamma tubulin. Both mNG-CCDC66 and mNG-CCDC66 (570-948) localized to the basal body, axoneme and the ciliary tip. In agreement with the strong centrosomal affinity of PACT, mNG-CCDC66-PACT localization was restricted to the centrosomes (Fig. 4A, S3B). We note that the localization profiles of these fusion proteins in CCDC66-depleted cells were similar in wild type cells, confirming that they are not due to overexpression artefacts (Fig. 1E, S3B, 4A).

**Figure 4.**
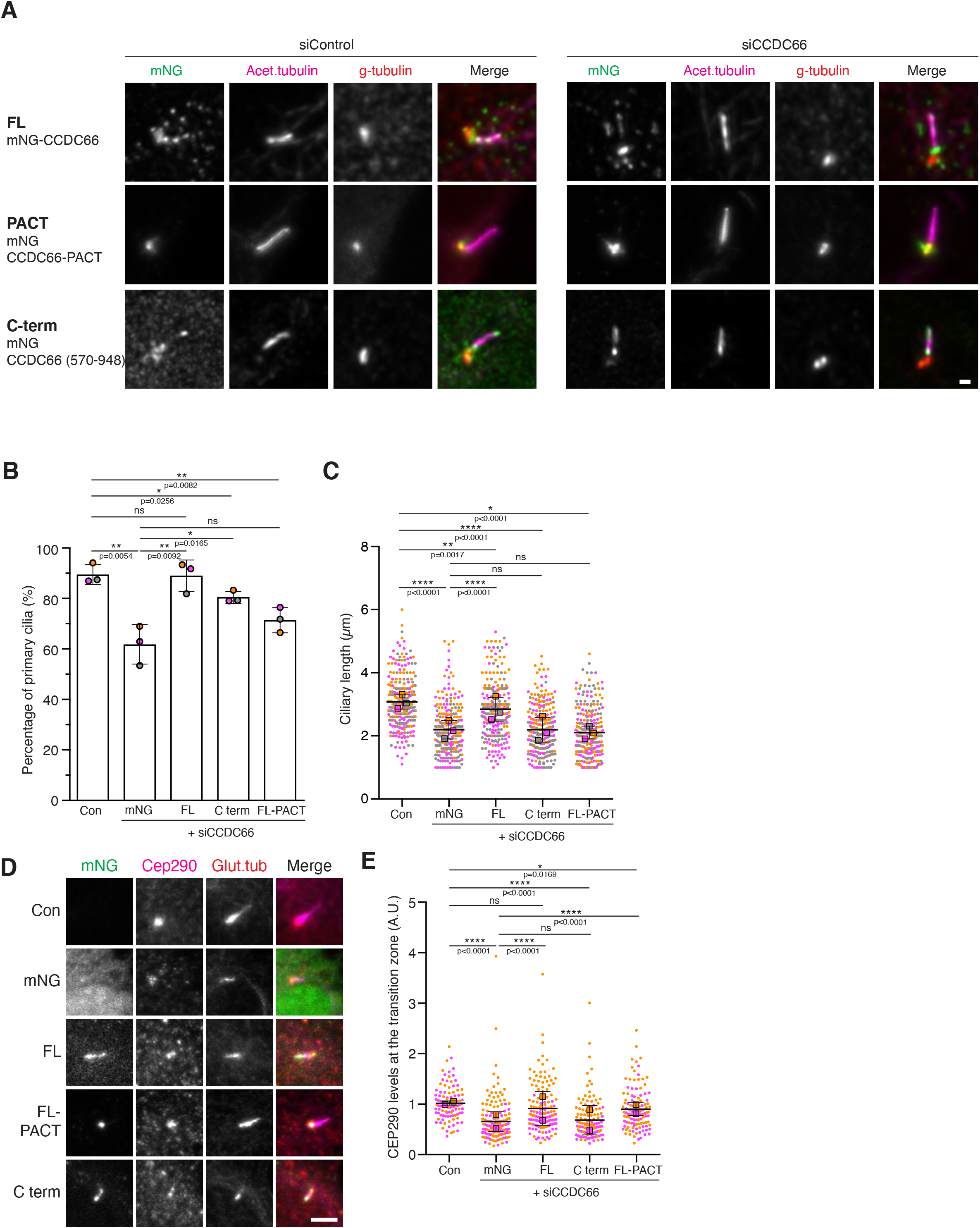
Axonemal and basal body pools of CCDC66 are required for cilium and transition zone assembly. **(A) Localization of CCDC66-PACT fusion and C terminal (570-948) fragment**. Control or CCDC66 depleted RPE1 cells stably expressing mNeonGreen-CCDC66 full length, CCDC66-PACT fusion and CCDC66 C-terminal (570-948) fragment were serum starved for 48 hours, fixed with 4% PFA and stained for mNeonGreen, acetylated tubulin and gamma tubulin. Scale bar: 1μm **(B-E) Rescue of phenotypes**. RPE1 cells with stably expressing mNeonGreen (mNG), mNeonGreen-CCDC66 full length (FL), mNeonGreen CCDC66-PACT fusion (FL-PACT) and CCDC66 C-terminal fragment (570-948) (C-term) were transfected with CCDC66 siRNA for two rounds and serum starved for 48 hours. As a control, RPE1 cells expressing mNeonGreen was transfected with control siRNA. **(A)** After fixation cells were stained for anti-acetylated tubulin for quantifying **(B)** percentage of cilium formation and **(C)** ciliary length, and **(D)** for mNeonGreen, CEP290 and polyglutamylated tubulin for quantifying CEP290 levels **(E)** at the transition zone. Data represent the mean ±SD. For cilium formation and cilia length quantifications, magenta, orange and grey represent individual values from three independent experiments (100 cells/experiment)For CEP290 levels, magenta and orange represent individual values from two independent experiments (50 cells/experiment, *P < 0.5, **P < 0.01, ***P < 0.001, ****P < 0.0001, ns: not significant one-way ANOVA).

Using these RPE1 stable lines, we performed rescue experiments for defective cilium assembly, axonemal elongation and targeting of CEP290 to the transition zone in CCDC66-depleted cells. mNG-CCDC66 expression rescued all three phenotypes to comparable levels to control siRNA-transfected mNG-expressing cells, demonstrating that these phenotypes are specific to CCDC66 depletion (Fig. 4B-E). mNG-CCDC66 (570-948) partially rescued the reduced ciliation defect (Fig. 4B), but not the shorter cilium length (Fig. 4C). Thus, the functional complexes formed by this fragment is not sufficient to rescue cilium length and is dependent on the activity of full-length protein. Finally, centrosomal CEP290 levels were partially restored by expression of mNG-CCDC66-PACT, but not mNG-CCDC66 (570-948), which might be because CCDC66-PACT created more binding sites for CEP290 at the centrosome (Fig 4D, 4E). This is consistent with the much higher centrosomal to cytoplasmic fluorescent intensity of mNG-CCDC66-PACT, indicating that the majority of cellular CCDC66 was sequestered at the centrosome (Fig. S3C). Given that mNG-CCDC66-PACT rescued only the defective transition zone recruitment of CEP290, these results further support that CCDC66 functions cilium assembly and axoneme elongation require cooperative activity of its distinct cellular pools at the microtubules, cilium and basal body.

### CCDC66 cooperates with the ciliopathy proteins CSPP1 and CEP104 during cilium length regulation

Little is known about how MAPs cooperate during cilium length regulation. To gain further insight into how CCDC66 regulates cilium length, we identified its proximity interaction partners in ciliated IMCD3 cells. We used IMCD3 cells as they ciliate with high efficiency and were used in proteomics studies of the primary cilium, which makes benchmarking easier (Mick et al. 2015; May et al. 2021). We generated IMCD3 cells stably expressing FLAG-miniTurbo-CCDC66 or FLAG-miniTurbo (control) using the Flip-IN system. As assessed by streptavidin staining, miniTurbo-CCDC66 biotinylated proteins at the centrosome and centriolar satellites in cycling cells and centriolar satellites, basal body, and cilium in serum-starved cells (Fig. S4A). Control cells expressing FLAG-miniTurbo exhibited biotinylation at the cytoplasm.

After validation of the cell lines, we performed large scale streptavidin pulldown of biotinylated proteins from cells serum starved for 48 h, analyzed them by mass spectrometry (Table 1) and defined high confidence CCDC66 interactome using Normalized Spectral Abundance Factor (NSAF) analysis (Firat-Karalar et al. 2014). The proximity interactome of ciliated cells consisted of 59 proteins. Analysis of this interactome by combining Gene Ontology (GO) analysis with literature mining revealed enrichment of centrosome/cilium/satellite proteins, MAPs, actin-binding proteins and proteins implicated in microtubule nucleation, which was visualized by Cytoscape (Fig. S4B). Notably, a number of these proximity interactors stand out by their known relationship to CCDC66 or involvement in cilium assembly and function. CSPP1, ARMC9, CEP104 and TOGARAM1 are axonemal proteins required for cilium length control (Das et al. 2015; Frikstad et al. 2019; Latour et al. 2020). Additionally, proteins mutated in retinal degeneration including Tubulin Tyrosine Ligase Like 5 (TTLL5), LCA5 and RPGRIP1L were also proximate to CCDC66 (den Hollander et al. 2007; Sergouniotis et al. 2014; Bedoni et al. 2016; Wiegering et al. 2018). Notably, the ciliated CCDC66 interactome did not include CEP290, which was previously identified as part of its interactome derived from asynchronous cells (Gupta et al. 2015; Conkar et al. 2017; Gheiratmand et al. 2019).

To elucidate the mechanisms by which CCDC66 regulates cilium length, we focused on the proximity interaction of CCDC66 with CSPP1 and CEP104. CSPP1 and CEP104 are MAPs that localize to the cilia and regulate axonemal length (Patzke et al. 2010; Frikstad et al. 2019; Yamazoe et al. 2020). Recent studies defined tubulin-binding TOG-domain containing proteins CEP104 and TOGARAM1 as potential axonemal polymerases (Yamazoe et al. 2020). However, the mechanisms by which their activity is regulated by other MAPs is not known (Farmer and Zanic 2021). To address this, we investigated the nature of relationship between CCDC66, CSPP1 and CEP104. First, we examined whether they physically interact or not by performing immunoprecipitation experiments in cells expressing GFP-CCDC66.

CSPP1 and CEP104 as well as known CCDC66 interactors PCM1 and CEP290 co-pelleted with GFP-CCDC66, but not the negative controls GFP or GFP-CCDC66 (1-408) (Fig. S4C). Notably, C-terminal 570-948 a.a. fragment did not interact with CSPP1, CEP104 and CEP290, which might explain why its expression did not restore cilium length defect (Fig S4C). Given that CCDC66 has multiple cellular pools, we next asked whether satellites and cytoplasmic microtubules are required for their interaction. As compared with control cells, loss of satellites by knocking out PCM1 compromised the interaction of GFP-CCDC66 with CSPP1 and CEP290, but not with CEP104 (Fig. 5AC). Consistent with this, CCDC66 co-localized with CEP290 at the centriolar satellites, but not at the transition zone (Fig. 1A). We note that CEP290 association defect might be due to reduced levels of total cellular CEP290 in PCM1^-/-^ cells (Fig. 5A). In contrast to loss of satellites, depolymerization of cytoplasmic microtubules by nocodazole treatment did not interfere with the ability of GFP-CCDC66 to interact with CSPP1 and CEP104 (Fig. S4D). Collectively, these results suggest that CCDC66 interacts with CSPP1 and CEP104 at the ciliary axoneme and that they might cooperate during cilium assembly and axoneme length regulation.

**Figure 5.**
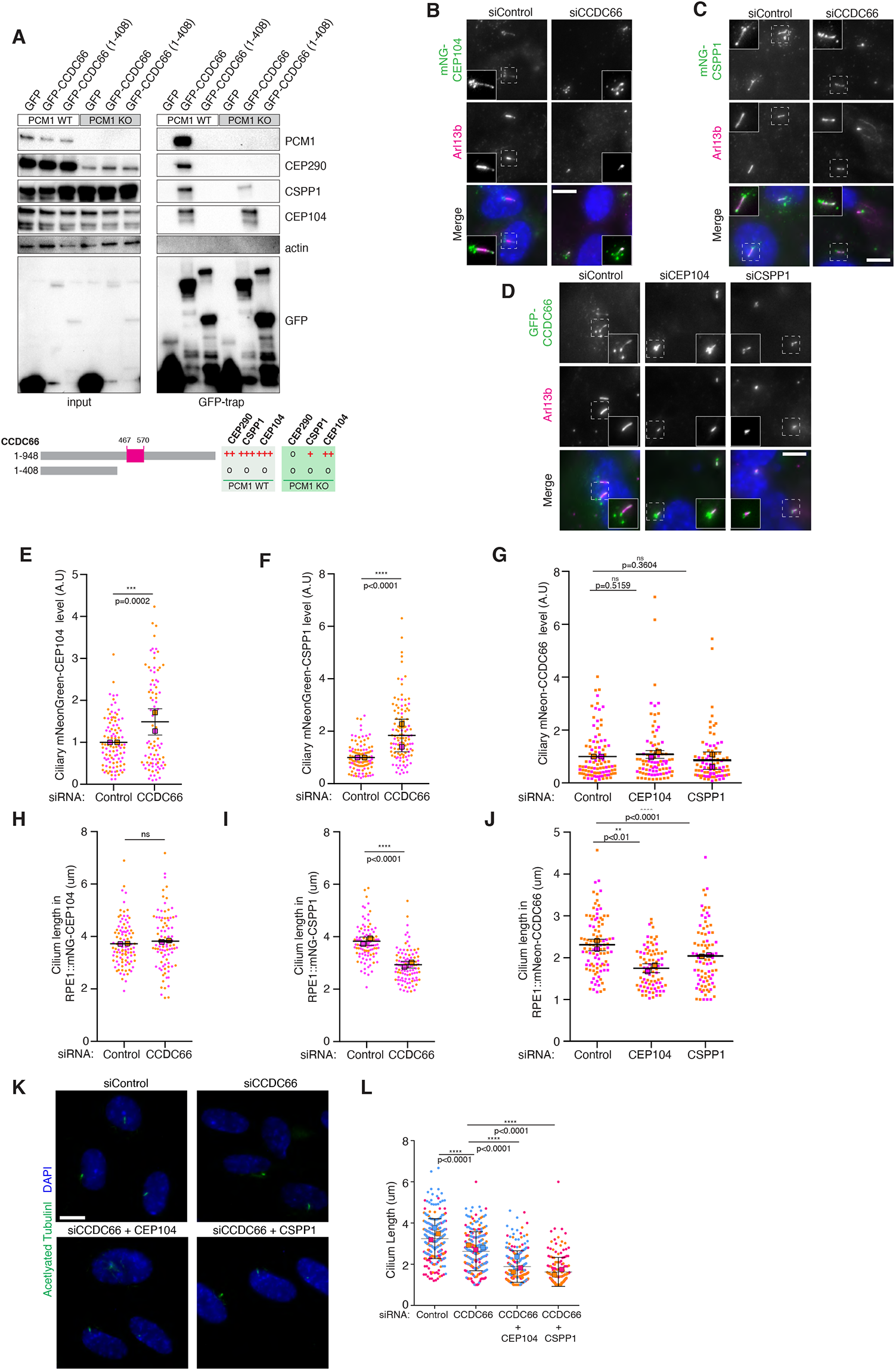
CCDC66 forms a functional complex with the ciliopathy proteins CSPP1 and CEP104. **(A) CCDC66 interacts with CSPP1 and CEP104**. PCM1 wild type (WT) and knockout (KO) HEK293T cells were transfected with EGFP, EGFP-CCDC66 full length and EGFP-CCDC66 (1-408) constructs. 2 days after transfection, cells were collected, lysed, and subjected to pull down with GFP Trap beads. Input and pellet were immunoblotted with anti GFP, PCM1, CEP290, CSPP1, CEP104 and actin as a control. The schematics summarizes the results of the pull down (**o**: no interaction, **+**: weak interaction, **++**: moderate interaction, **+++**: strong interaction). **(B-C, E-F and H-I) CCDC66 negatively regulates ciliary CSPP1 and CEP104 level. (B)** RPE1::mNG-CEP104 or **(C)** RPE1::mNG-CSPP1 cells were transfected with two rounds of control or CCDC66 siRNA and serum starved for 48 hours. Following fixation, cells were stained for anti mNeonGreen and ARL13B antibodies, and DAPI for visualization of DNA. **(E)** Ciliary mNG-CEP104 or **(F)** mNG-CSPP1 signal was quantified by measuring the mNeon signal intensity using the area covered by ARL13B signal, subtracting the background signal, and dividing it by the length of the cilia. Cilium length **(H-I)** were measured and plotted. Data represent the mean ±SD. Magenta and orange represent individual values from two independent experiments. (50 cilia/experiment, *P < 0.5, **P < 0.01, ***P < 0.001, ****P < 0.0001, ns: not significant t-test) Scale bar: 10μm **(D-G-J) Regulation of CCDC66 ciliary targeting by CEP104 and CSPP1**. RPE1::mNeon-CCDC66 cells were transfected with two rounds of control, CEP104 or CSPP1 siRNA and serum starved for 48 hours. **(D)** Following fixation, cells were stained for anti-mNeon and ARL13B antibodies, and DAPI for visualization of DNA. **(G)** Ciliary mNeon-CCDC66 signal was quantified by measuring the mNeon signal intensity using the area covered by ARL13B signal and dividing it by the length of the cilia. Cilium length **(J)** were plotted. Data represent the mean ±SD. Magenta and orange represent individual values from two independent experiments. (50 cilia/experiment, *P < 0.5, **P < 0.01, ***P < 0.001, ****P < 0.0001, ns: not significant t-test) Scale bar: 10μm **(K-L) Co-depletions of CEP104 or CSPP1 along with CCDC66 results shorter cilia than only CCDC66. (K)** RPE1 cells were transfected with two rounds of control, CCDC66, CCDC66 along with CEP104 and CCDC66 along with CSPP1 siRNA and fixed at 48 hours serum starvation. Following fixation, cells were stained for antiAcetlyated tubulin antibody and DAPI. (L) Cilium length is plotted. Magenta, orange and cyan represent individual values from three independent experiments. Error bars represent ±SD. Square boxes represents the mean value of each experiment for each group. (50 cilia/experiment, *P < 0.5, **P < 0.01, ***P < 0.001, ****P < 0.0001, ns: not significant one-way ANOVA) Scale Bar: 10μm

Next, we asked whether CCDC66, CEP104 and CSPP1 depend on each other for localization to the primary cilium using previously characterized RPE1 cells stably expressing their fluorescent fusions (Conkar et al. 2017; Frikstad et al. 2019). We depleted CEP104 and CSPP1 using previously described siRNAs, which we validated by quantification of ciliogenesis efficiency and cilium length (Fig. S4E-G) (Patzke et al. 2010; Yamazoe et al. 2020). Ciliary level of mNG-CSPP1 and mNG-CEP104 increased in CCDC66-depleted cells relative to control cells, identifying an inhibitory role for CCDC66 in their ciliary recruitment (Fig. 5B, 5C, 5E, 5F). CEP104 and CSPP1 depletions did not change in the ciliary level of mNeon-CCDC66 (Fig. 5D-G).

In addition ot their ciliary localization dependency, we investigated the functional dependency of CCDC66, CSPP1 and CEP104 during axoneme elongation. To this end, we first examined whether stable expression of mNG-CEP104 and mNG-CSPP1 restores the cilium length defect associated with CCDC66 depletion. RPE1::mNG-CEP104 or RPE1::mNG-CSPP1 cells were transfected with control or CCDC66 siRNAs and serum starved for 48 h. Intriguingly, expression of mNG-CEP104, but not mNG-CSPP1, compensated for the cilium length defects of CCDC66-depleted cells (Fig. 5H, 5I). Given the role of TOG domain of CEP104 in microtubule polymerization, this result suggests that CCDC66 might be involved in cilium length control via regulating CEP104-mediated axonemal polymerization (Yamazoe et al. 2020). Consistently, depletion of CEP104 by RNAi in RPE1::mNG-CCDC66 cells further reduced cilium length (Fig.5J). Next, we investigated whether CSPP1 or CEP104 co-depletion further exacerbates cilium length defect associated with CCDC66 depletion. In both cases, the cilia that formed in co-depleted cells were shorter relative to cells depleted for only CCDC66, indicating that these proteins are not functionally redundant (Fig. 5K-L). Taken together, our findings suggest that CCDC66 and CSPP1 functions during cilium length control in part via regulating CEP104-mediated axonemal microtubule polymerization.

### CCDC66 depletion perturbs cilium content regulation and Hedgehog pathway response

To determine how ciliary defects associated with CCDC66 loss affects primary cilium function, we characterized the cilia that forms in CCDC66-depleted cells for their content and response to Hedgehog pathway stimuli. First, we examined the constitutive ciliary membrane proteins implicated in ciliary signaling, including the small GTPase ARL13B and ciliary GPCR protein SSTR3. Ciliary ARL13B levels were similar between control and CCDC66-depleted cells (Fig. 6A, B). However, there was about 1.3-fold increase in somatostatin receptor SSTR3 ciliary level upon CCDC66 depletion (Fig. 6C, D). Next, we examined Hedgehog pathway activation in RPE1 cells transfected with control and CCDC66 siRNA. Upon stimulation of cells with Hedgehog ligands at the cilium, the GPCR SMO enters the cilium and GPR161 moves out of the cilium, which eventually leads to transcriptional activation of Hedgehog target genes (Mukhopadhyay and Rohatgi 2014). As functional readouts for Hedgehog pathway activation, we quantified the efficiency of ciliary entry of SMO and Gli1 upregulation (Fig. 6E-J). To this end, ciliated control and CCDC66 depleted cells were treated with 100 nM Smoothened agonist (SAG) for 24 h and processed by immunofluorescence or quantitative PCR. In both control and CCDC66-depleted cells, SMO did not localize to cilia under basal conditions (Fig. 6E). Upon SAG stimulation, the ciliary level of SMO increased about 3-fold in control cells and 2.3-fold in CCDC66-depleted cells (Fig. 6E, F). This significant reduction in ciliary SMO enrichment shows that CCDC66 loss interferes with its SAG-induced ciliary accumulation. Additionally, we compared the ciliary distribution of SMO upon SAG treatment and found that the fraction of cells with “tip” localization of SMO as well as its levels at the ciliary tip were reduced upon CCDC66 depletion (Fig. 6G-I). Finally, we examined the downstream consequences of these alterations by quantification of Gli1 transcriptional upregulation in SAG-treated cells. In contrast to control cells, CCDC66 depleted cells failed to upregulate Gli1 in response to SAG treatment (Fig. 6J). Collectively, these findings demonstrate that CCDC66 is required for the formation of Hedgehog-competent cilia.

**Figure 6.**
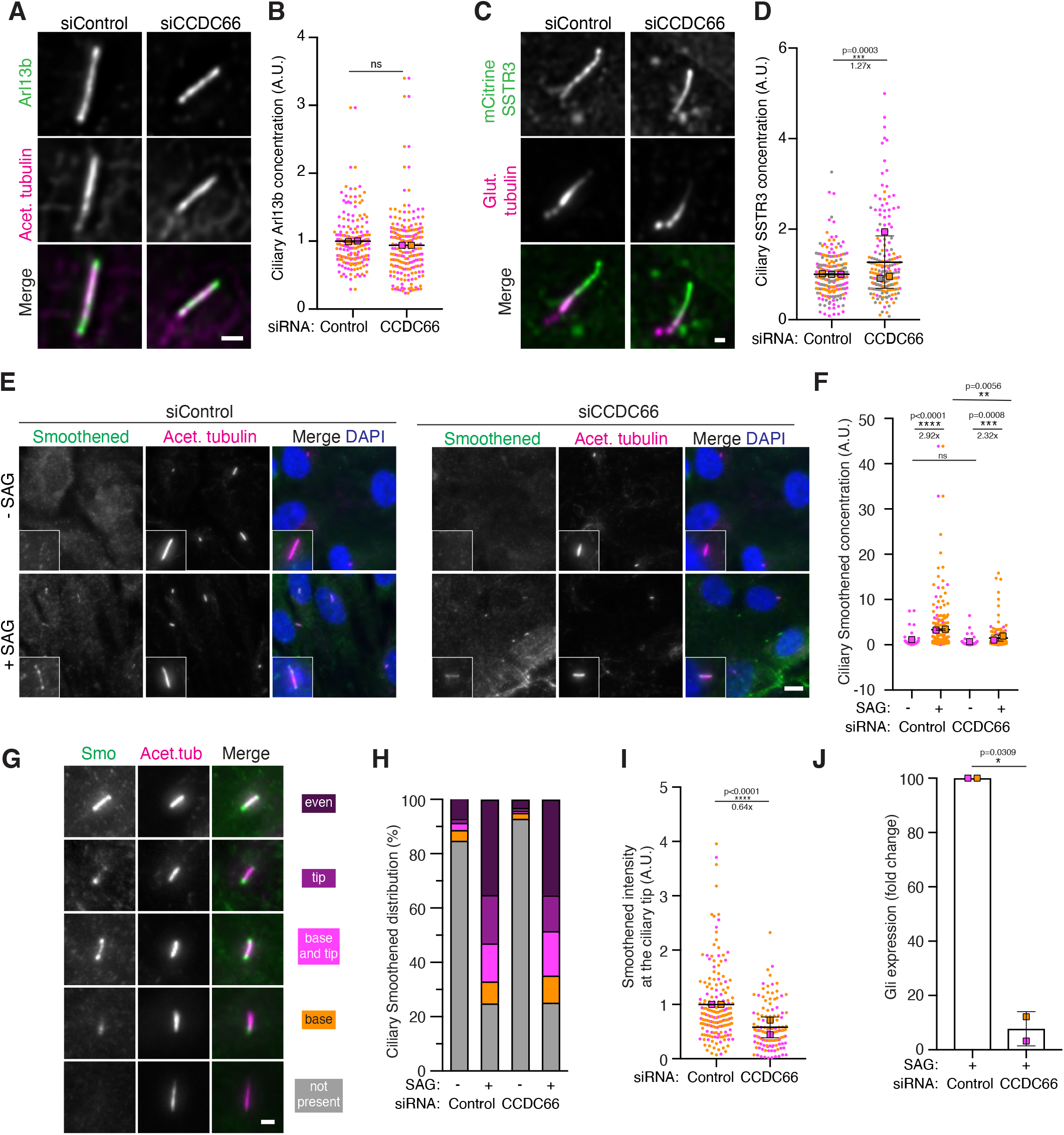
CCDC66 impairs Hedgehog pathway activation. **(A-B) CCDC66 is not required for ciliary ARL13B recruitment**. RPE1 cells were transfected with two rounds of control or CCDC66 siRNA and serum starved for 48 hours. Cells were fixed and stained for ARL13B and acetylated tubulin antibodies. Ciliary ARL13B levels were quantified by measuring the ARL13B intensity at cilia, subtracting the background signal, multiplied by the signal area, and dividing by the cilium length Data represent the mean ±SD. Magenta and orange represent individual values from two independent experiments. (50 cilia/experiment, ns: not significant t-test) Scale Bar: 1μm **(C-D) CCDC66 depletion causes ciliary SSTR3 accumulation**. RPE1::mCitrine-SSTR3 cells were transfected with two rounds of control or CCDC66 siRNA and serum starved for 48 hours. Cells were fixed with 4%PFA and stained for anti-polyglutamylated tubulin. Ciliary mCitrine-SSTR3 levels were quantified by measuring the SSTR3 intensity at cilia, subtracting the background signal, multiplied by the signal area, and dividing by the cilium length Data represent the mean ±SD. Magenta and orange represent individual values from two independent experiments. (50 cilia/experiment, ***P < 0.001 t-test) Scale bar: 1μm **(E-I) CCDC66 depletion compromises ciliary recruitment and distribution of SMO**. RPE1 cells were transfected with two rounds of control or CCDC66 siRNA and serum starved for 24 hours and incubated with (0.01%) DMSO or 100 nM SAG for the subsequent 24 hours. **(E)** Cells were fixed and stained for anti-Smoothened, anti-acetylated tubulin and DAPI. Hedgehog activation was assessed by **(F)** ciliary Smoothened levels were quantified by measuring the SMO intensity at cilia, **(G-H)** assessment of the localization of Smoothened in the absence and presence of SAG, and **(I)** the ciliary tip levels of Smoothened in SAG treated cells. Data represent the mean ±SD.. Magenta and orange represent individual values from two independent experiments. (50 cilia/experiment, *P < 0.5, **P < 0.01, ***P < 0.001, ****P < 0.0001, ns: not significant, two way ANOVA for F, t-test for I) Scale bar: 10-μm for C and 2μm for E. **(J) CCDC66 depletion affects GLI1 upregulation as a response of Hedgehog signaling activation**. Upregulation of GLI1 expression in control and CCDC66 depleted cells treated with (0.01%) DMSO or 100mM SAG. GLI1 mRNA was quantified by qPCR before SAG treatment and 24 h after SAG treatment, and its fold change is normalized to control cells (=100). Results shown are the mean of two independent experiments (±SD) (*P < 0.5, t-test)

## Discussion

CCDC66 is a MAP that localizes to the centriolar satellites, the centrosome and the primary cilium and is implicated in ciliopathies affecting retina and brain. The results of our study reveal two specific functions of CCDC66 at the primary cilium. First, it is required for assembling the primary cilium with high efficiency and at the right composition. In particular, it functions directly or indirectly during Hedgehog pathway activation. It mediates these functions in part by regulating transition zone assembly and basal body recruitment of the IFT-B complex. Second, CCDC66 interacts with the ciliopathy proteins CEP104 and CSPP1 and controls axonemal length. The results of our study have important implications for uncovering the mechanisms that underlie the structural and functional complexities of the primary cilium as well as pathogenesis of CCDC66-linked ciliopathies.

The distinct localization profiles of CCDC66 full length and truncation constructs allowed us, for the first time, to address important questions regarding the significance of functional and dynamic compartmentalization within the centrosome/cilium complex. For example, primary cilium assembly and disassembly dynamics have so far been studied by monitoring dynamic behavior of ciliary membrane proteins (Westlake et al. 2011; Lu et al. 2015; Jewett et al. 2021; Quanlong et al. 2021). Our analysis of the localization dynamics of CCDC66 during ciliogenesis is the first report of the kinetics of the ciliary axoneme in mammalian cells. Another key finding of our study is that we identified differences in CCDC66 interaction partners at different cellular locations: a satellite-dependent interaction with CEP290 and a satellite-independent interaction with CEP104, which is suggestive of different mechanisms by which CCDC66 mediates its functions at the centriolar satellites versus the primary cilium. Future studies are required to investigate the molecular basis underlying these differences.

Formation of signaling-competent cilia is a multistep process that requires nucleation of basal body-templated axonemal microtubules and their elongation to and maintenance at a steady-state length. Our findings provide important insight into how the microtubule-based core of the cilium is assembled and maintained by the coordinated activity of MAPs and ciliary signaling/transport proteins. We showed that CCDC66 binds to microtubules, localizes to the axoneme and the cilium tip and interacts with axonemal/tip proteins CEP104 and CSPP1. CEP104 contains the evolutionarily conserved tubulin-binding TOG domains (Farmer and Zanic 2021). Given the reported functions of TOG-array proteins in microtubule dynamics such as microtubule polymerization, CEP104 has been proposed to regulate polymerization of axonemal microtubules (Das et al. 2015; Yamazoe et al. 2020). The cilium length defect of CCDC66-depleted cells is compensated by expression of mNG-CEP104, suggesting that CCDC66 regulates CEP104-mediated axonemal microtubule polymerization. Future *in vitro* studies will be critical in informing on how CCDC66-associated ciliary complexes regulate microtubule dynamics.

Centriolar satellites have emerged as regulators of cilium assembly and composition in part via centrosomal and ciliary targeting of proteins implicated in these processes (Gheiratmand et al. 2019; Quarantotti et al. 2019; Aydin et al. 2020). During the live imaging experiments, we observed the centriolar satellite pool of CCDC66 disappearing as the ciliary pool of CCDC66 appeared as the cilium assembled. Considering the structural and functional defects observed in the absence of CCDC66, this observation places CCDC66 as a regulatory factor in primary cilium assembly. Of course, whether the source of ciliary CCDC66 is newly synthesized protein or CCDC66 translocating from centriolar satellites remains to be answered. Our results on how CCDC66 regulates cilium content provide a model for how centriolar satellites achieve this function. We show that CCDC66 is required for transition zone assembly via regulating ciliary targeting of CEP290. Importantly, CCDC66 does not localize to the transition zone, instead interacting with CEP290 at the centriolar satellites, suggesting that the satellite granules function as scaffolds for mediating this interaction. Taking into account previous work that revealed a critical role for the satellite pool of CEP290 during transition zone assembly, we propose the following model for the functional significance of the CCDC66-CEP290 interaction (Kobayashi et al. 2014; Tu et al. 2018): CCDC66 maintains the satellite pool of CEP290 by tethering it to PCM1, and satellites regulate basal body targeting of CEP290 by sequestration or active transport. This model is supported by two lines of evidence. First, we found an inverse correlation of the number and integrity of satellite granules with the growth of primary cilium. Second, tethering CCDC66 to the centrosome via its PACT fusion is accompanied by lack of its satellite pool and restoration of CEP290 levels at the transition zone in CCDC66-depleted cells. In addition to CEP290, the centriolar satellite pool of CCDC66 might also regulate cilium content via ciliary targeting of the transport complexes. Consistently, CCDC66 depletion impaired ciliary localization of several components of the BBSome complex, which was identified as part of the centriolar satellite interactome (Conkar et al. 2017; Gheiratmand et al. 2019; Quarantotti et al. 2019). Whether and if so how CEP290 and ciliary transport proteins exchange between satellites and cilium needs to be further investigated.

We identified the *in vivo* proximity interactome of CCDC66 in ciliated cells. To this end, we used confluent IMCD3 cultures serum starved for 48 h, which contain about 90% ciliated cells. While the resulting map provided insight into ciliary functions and mechanisms of CCDC66, we note that it has low overlap with the published proximity interactomes of the cilium generated by APEX-fusions of ciliary targeting sequences of membrane proteins (Mick et al. 2015; Kohli et al. 2017; May et al. 2021). There are two possible explanations for the low overlap. First, CCDC66 localizes to the centriolar satellites, basal body and primary cilium in ciliated cells. Therefore, its ciliary interactions might be of lower abundance than that of its other cellular pools. Second, CCDC66 might mediate its structural functions at the axoneme by forming a stable complex, which would limit its interactions with other ciliary proteins. This is supported by the FRAP data that revealed that ciliary pools of CCDC66 and its interactor CSPP1 are immobile (Conkar et al. 2019; Frikstad et al. 2019). Future studies that specifically identify its ciliary interactions are required to distinguish between these possibilities.

In summary, our findings identify a complex molecular and functional relationship between the different compartments of the centrosome/cilium complex and provide directions for future studies on the molecular basis of differential complex formation and intricate interplay of ciliopathy proteins during cilium biogenesis and function. Future studies on elucidating the functions of CCDC66 in different cell types and tissues will contribute to our understanding of how its deregulation cause ciliopathies affecting eye and brain.

## Materials and Methods

### Cell culture, transfection and lentiviral transduction

Human embryonic kidney (HEK293T, ATCC, CRL-3216) cells were cultured with Dulbecco’s Modified Eagle’s Medium DMEM (Pan Biotech, Cat. # P04-03590) supplemented with 10% Fetal Bovine Serum (FBS, Life Technologies, Ref. # 10270-106, Lot # 42Q5283K) and 1% penicillin-streptomycin (Gibco, Cat. # 1540-122). Human telomerase immortalized retinal pigment epithelium cells (hTERT-RPE, ATC, CRL-4000) and mouse kidney medulla collecting duct cells IMCD3:Flip-In cells were cultured with DMEM/F12 50/50 medium (Pan Biotech, Cat. # P04-41250), supplemented with 10% FBS and 1% penicillin-streptomycin. RPE1::mNG-CEP104 and RPE1::mNG-CSPP1 cell lines were described previously (Frikstad et al. 2019). All cell lines were tested for mycoplasma by MycoAlert Mycoplasma Detection Kit (Lonza). RPE1 cells were transfected using Lipofectamine LTX (Thermo Fisher). HEK293T cells were transfected with the indicated plasmids using 1 μg/μl polyethylenimine, MW 25 kDa (PEI, Sigma-Aldrich, St. Louis, MO).

Lentivirus were generated using pcDH-mNG-CCDC66, pcDH-mNG-CCDC66 (570-948), pcDH-mNG-CCDC66-PACT and pCDH-EF1-mNeonGreen-T2A-Puro, and pLVPT2-mScarlet-Arl13b plasmids as transfer vectors. RPE1 cells were transduced with the indicated lentivirus and selected with 6 μg/ml puromycin for four to six days until all the control cells died. IMCD3 cells stably expressing FLAG-miniTurbo and FLAG-miniTurbo-CCDC66 were generated using previously described protocols. Briefly, cells were co-transfected with pcDNA5.1-FRT/TO expression vectors and pOG44 at a ratio of 1:7 using Lipofectamine LTX, selected with 300 μg/ml hygromycin B and individual colonies were picked and validated by immunofluorescence.

For cilium assembly experiments, cells were washed twice with PBS and incubated with DMEM/F12 50/50 supplemented with 0.5% FBS and 1% penicillin-streptomycin for the indicated times. For cilium disassembly experiments, cells that were incubated with 0.5% FBS for 48 hours were washed twice with PBS and incubated with DMEM/F12 50/50 supplemented with 10% FBS and 1% penicillin-streptomycin for the indicated times. For Hedgehog pathway activation, cells were incubated with 100 nm Smoothened agonist (SAG, EMD Millipore) or DMSO for 24 h following 24 hours of serum starvation.

### Plasmids and siRNAs transfections

pDEST-GFP-CCDC66, pDEST-GFP-CCDC66^RR^ and pDEST-Flag-CCDC66 plasmids were previously described (Conkar et al. 2017). Full-length CCDC66 was cloned into pcDNA5.1-FRT/TO-FLAG-miniTurbo vector to generate IMCD3 stable lines using the Flip-In approach. Full-length CCDC66, CCDC66 (570-948) and CCDC66-PACT were cloned into pCDH-EF1-mNeonGreen-T2A-Puro lentiviral expression plasmid. siRNA resistant mNeonGreen-CCDC66 was amplified from siRNA resistant GFP-CCDC66^RR^ plasmid and cloned into pCDH-EF1-mNeonGreen-T2A-Puro plasmid. CCDC66 was depleted using an siRNA with the sequence 5′-CAGTGTAATCAGTTCACAAtt-3′ (Thermo Scientific). Silencer Select Negative Control No. 1 (Thermo Scientific) was used as a control. CSPP1 and CEP104 were depleted by RNAi using previously described siRNAs (Patzke et al. 2010; Yamazoe et al. 2020). TOGARAM1 was depleted using siRNA with sequence 5′-CCUCGUAAUUCCUUAGAAA-3′ (Thermo Scientific) Cells were seeded onto coverslips at 70% confluency and transfected with 50 nM of siRNA in two sequential transfections using Lipofectamine RNAiMAX (Life Technologies, Ref. # 13778-150, Lot # 2009103) in OPTI-MEM (Life Technologies, Cat. # 31985062) according to the manufacturer’s instructions. Depletion of proteins was confirmed at 48 h or 72 h after transfection by immunofluorescence and immunoblotting.

### Immunofluorescence and antibodies

Cells were grown on coverslips, washed twice with PBS and fixed with either ice cold methanol at −20°C for 10 minutes or 4% PFA in cytoskeletal buffer (10 mM PIPES, 3 mM MgCl^2^, 100 mM NaCl, 300 mM sucrose, pH 6.9) supplemented with 5 mM EGTA and 0.1% Triton X for 15 minutes at 37°C. After washing twice with PBS, cells were blocked with 3% BSA (Capricorn Scientific, Cat. # BSA-1T) in PBS + 0.1% Triton X-100 and incubated with primary antibodies in blocking solution for 1 hour at room temperature. Cells were washed three times with PBS and incubated with secondary antibodies and DAPI (Thermo Scientific, cat# D1306) at 1:2000 for 45 minutes at room temperature. Following three washes with PBS, cells were mounted using Mowiol mounting medium containing N-propyl gallate (Sigma-Aldrich).

Detergent incubation to assess axonemal association of CCDC66 is adapted from (Nachury et al. 2007). Briefly, cells were washed twice with PHEM buffer (60 mM PIPES, 25 mM HEPES, 10 mM EGTA, 4 mM MgSO_4_, pH 7.0) and incubated with either PHEM or PHEM+ 0.5% Triton-X for 30 seconds. After incubation, they were fixed with 4% PFA at 37°C and processed for microscopic analysis. Primary antibodies used for immunofluorescence were rabbit anti-IFT88 (13967-1-AP, Proteintech) at 1:50, rabbit anti-IFT81 (11744-1-AP, Proteintech) at 1:50, rabbit anti-CP110 (A301-344A, Betyl) at 1:500, mouse anti-PCM1 1:1000 (sc-398365, Santa Cruz Biotechnology), rabbit anti-CEP290 (800 338 9579, Betyl) at 1:500, mouse anti-BBS1 (sc-365138, Santa Cruz Biotechnology) at 1:50, mouse anti-BBS2 (sc-365355, Santa Cruz Biotechnology) at 1:50, mouse anti-BBS3 (sc-390021, Santa Cruz Biotechnology) at 1:50, mouse anti-CEP164 (sc-515403, Santa Cruz Biotechnology) at 1:1000, mouse anti-Smoothened (sc-166685, Santa Cruz Biotechnology) at 1:50, mouse anti-polyglutamylated tubulin (AG-20B-0020, clone GT335, Adipogen), mouse anti-gamma tubulin (T5326, clone GTU-88, Sigma) at 1:1000, mouse anti-acetylated tubulin (clone 6-11B, 32270, Thermo Fischer) at 1:10000), rabbit anti-ARL13B (17711-1-AP, Proteintech) at 1:50, mouse anti-ARL13B (75-287, NIH Neuromab) at 1:100, mouse anti-GFP (3E6, Invitrogen) at 1:750. Rabbit anti-GFP and rabbit anti-PCM1 antibodies were generated and used for immunofluorescence as previously described (Conkar et al. 2017). Anti-CCDC66 antibody was generated by immunizing rats (Koc University, Animal Facility) with His-MBP-tagged mouse CCDC66 (clone 30626499) comprising amino acids 1-756 purified from Hi5 insect cells. The antibody was affinity purified against His-MBP-mCCDC66 (aa 1-756) and used at 0.5 ug/ml for immunofluorescence. Secondary antibodies used for immunofluorescence experiments were AlexaFluor 488-, 568- or 633-coupled (Life Technologies) and they were used at 1:2000. Secondary antibodies used for Western blotting experiments were IRDye 680-and IRDye 800-coupled and were used at 1:15,000 (LI-COR Biosciences), Peroxidase AffiniPure Donkey Anti-Mouse IgG (H+L) (715-035-150, Jackson ImmunoResearch) and Peroxidase AffiniPure Goat Anti-Rabbit IgG (H+L) (111-035-144, Jackson ImmunoResearch).

### Microscopy and image analysis

Time lapse live imaging was performed in an incubation chamber on Leica SP8 confocal microscope with HC PL APO CS2 63x 1.4 NA oil objective. For imaging CCDC66 localization dynamics during cilium assembly, cells were incubated with 0.5% FBS in DMEM-12 after 2 rounds of siRNA transfection and imaged overnight at every 12 minutes per frame in 512×512 pixel format. For cilium disassembly, cells that were transfected with siRNA, serum starved for 2 days. Following serum stimulation, cilium disassembly was imaged overnight at every 12 minutes in 512×512 pixel format. For protein level and localization percentage quantifications, images were acquired with Leica DMi8 fluorescent microscope with a stack size of 8 μm and step size of 0.3 μm in 1024×1024 format using HC PL APO CS2 63x 1.4 NA oil objective. Higher resolution images were taken by using HC PL APO CS2 63x 1.4 NA oil objective with Leica SP8 confocal microscope. Images were processed using Image J (National Institutes of Health, Bethesda, MD).

Quantitative immunofluorescence of centrosomal and ciliary levels of proteins was performed by acquiring a z-stack of cells using identical gain and exposure settings, determined by adjusting settings based on the fluorescence signal in the control cells. The z-stacks were used to assemble maximum-intensity projections. The centrosome regions in these images were defined by centrosomal marker staining for each cell, and the total pixel intensity of a circular 2.5 μm^2^ area centered on the centrosome in each cell was measured using ImageJ and defined as the centrosomal intensity. For transition zone quantification, 2.5 μm^2^ area above basal body is measured. Basal body is determined glutamylated tubulin signal. The ciliary regions in these images were defined by ARL13B or acetylated tubulin for each cell. For the basal body acetylated tubulin levels, 2.2 μm^2^ region of interest (ROI) was drawn and 3 random areas were quantified using this ROI. The background intensity was subtracted from their average. Background subtraction was performed by quantifying fluorescence intensity of a region of equal dimensions in the area neighboring the centrosome or cilium. Ciliary protein levels were determined by dividing fluorescence signal of the protein to the cilium length, which was quantified using ARL13B or acetylated tubulin staining Centrosomal and ciliary. Protein levels were normalized relative to the mean of control group.(=1). Line analysis was performed by drawing a line covering the basal body and ciliary signal of GFP-CCDC66 signal and the marker. Each protein signal is normalized to average intensity of the it’s own signal and plotted. Primary cilium formation was assessed by counting the total number of cells and the number of cells with primary cilia, as detected by Arl13b or acetylated tubulin staining and DAPI staining. Localization percentage quantifications were performed by counting the number of cilia using a ciliary marker such as acetylated tubulin or ARL13B and the number of cilia with the desired protein signal and calculating the percentage of positivity for the corresponding protein. The width of the cilium from U-ExM experiments were quantified as the width of the area where Arl13b signal starts to appear on cilium is measured.

To assess Hedgehog pathway activation, ciliary Smoothened level was measured by the background subtracted ciliary Smoothened signal divided by the ciliary length. Ciliary tip levels of Smoothened was quantified by measuring Smoothened signal with 0.5 μm^2^ ROI above the acetylated tubulin marker and subtracting it from the background signal. Ciliary Smoothened distribution categories were determined according to the observed Smoothened distribution patterns. All data acquisition was done in a blinded manner.

For analysis of live imaging movies, the frame in which cilium has started to form was determined by the bulging of CCDC66 or Smoothened signal from the basal body. Quantification of ectocytosis, breakage (scission) and ripping off events was done manually by inspection of each cilium. Breakage events were distinguished from ectocytosis events by the length of the ciliary piece released (ectocytosis event < 0.5 μm, breakage/rip-off event > 0.5 μm). All values representing levels were normalized relative to the mean of control group. (=1). Statistical significance was determined by Student’s t-test using Prism (GraphPad, La Jolla, CA).

### Cell lysis and immunoblotting

Cells were washed with PBS twice and lysed in the lysis buffer (50 mM Tris pH 7.6, 150 mM NaCl, 1% Triton X-100), tumbled at 4°C for 40 minutes and centrifuged at 13.000 R.P.M. Protein concentration was measured with Bradford solution (Bio-Rad Laboratories, CA, USA). The resulting supernatant was added with 6x sample buffer, boiled for 10 minutes at 95°C. Proteins were separated by SDS-PAGE, transferred to nitrocellulose membranes (Bio-Rad Laboratories, CA, USA) and blocked with 5% milk in TBS with 1%Tween-20. Primary antibody incubation was performed at 4°C overnight or at room temperature for 2 hours. Secondary antibody incubation was performed at room temperature for 1 hour. Membranes were washed with PBS for 15 minutes and scanned in Li-Cor Odyssey® Infrared Imaging System software (Li-Cor Biosciences) at 169 μm. ChemiDoc MP Imaging System (Bio-Rad Laboratories, CA, USA) was used for peroxidase coupled secondary antibodies. SuperSignal™ West Pico PLUS Chemiluminescent Substrate (34577, Thermo Scientific),SuperSignal™ West Dura Extended Duration Substrate (37071Thermo Scientific),SuperSignal™ West Femto Maximum Sensitivity Substrate (34094, Thermo Scientific) were used as chemiluminescence reagents. Quantifications of band intensities and cropping of the images were performed in ImageJ.

### Biotin identification with miniTurbo and mass spectrometry analysis

IMCD3 cells stably expressing miniTurbo or miniTurbo-CCDC66 were used. For mass spectrometry analysis, each cell type was grown in 5×15 cm plates in DMEM/F12 medium supplied with 10% FBS and 1% penicillin-streptomycin. The ciliated cell populations were generated after growing cells to 100% confluency and serum starving them for 48 hours in DMEM/F12 with 0% FBS. Both cell populations were incubated with 500 μM biotin for 30 minutes. After washing two times with PBS at room temperature, cells were collected and lysed in RIPA buffer (50 mM Tris pH 8.0, 150 mM NaCl, 0.1% SDS, 0.5% sodium deoxycholate, 1% Triton X-100) freshly supplied with 1x ProBlock protease inhibitor cocktail (GoldBio), and 1 mM Phenylmethylsulfonyl fluoride (PMSF). Cell lysates were sonicated, and their protein concentration were determined. 2.5% of the lysate was saved as the initial sample. Lysates were centrifuged at 14.000 R.P.M. for 1 hour at 4°C. Pellet and 50 μl of supernatant was saved for SDS-PAGE analysis. Remaining supernatant was incubated with 200 μl Streptavidin agarose beads (Thermo Scientific) for 16 hours at 4°C. Following incubation, beads were washed twice with RIPA buffer, once with 1 M KCl, once with 0.1 M Na_2_CO_3_, once with 2 M urea in 10 mM Tris-HCl pH 8.0, and lastly two times with RIPA buffer. For mass spectrometry analysis, beads were resuspended in 100 μl of 50 mM ammonium bicarbonate and performed at KUPAM proteomics facility as previously described (Gurkaslar et al. 2020). For miniTurbo experiment data presented in the table or network format in Fig. S4 and Table 1, data were derived from two biological replicates and two technical replicates.

For mass spectrometry analysis, Normalized Spectral Abundance Factor (NSAF) values were generated for each protein by dividing each Peptide Spectrum Match (PSM) value by the total PSM count in that dataset. Datasets were filtered as follows: First, proteins that were present only in the control dataset and in only one of the technical datasets were removed. NSAF values in the CCDC66 dataset were divided by the corresponding NSAF value from the control dataset to calculate an enrichment score for filtering proteins that were more abundant in the cycling or ciliated datasets. For proteins present in both experimental replicas, the average of enrichment score is calculated. Proteins with enrichment score <2 were removed. Next, the remaining proteins were submitted to CRAPome (https://reprint-apms.org), which is a contaminant repository for mass spectrometry data collected form affinity purification experiments and a list with contaminancy percentage (%) was calculated (Mellacheruvu et al. 2013). Proteins with contaminancy percentage more than 25% were considered as a contaminant and removed. This cut-off value was chosen depending on whether there was a known interaction partner of CCDC66 within the range of that value. Lastly, proteins were listed according to their NSAF values and the interactome map was generated using Cytoscape (Shannon et al. 2003).

### RNA isolation, cDNA Synthesis, and qPCR

Total RNA was isolated from control and IMCD PCM1 KO cells before SAG treatment and 24 h after SAG treatment using NucleoSpin RNA kit (Macherey-Nagel) according to the manufacturer’s protocol. Quantity and purity of RNA were determined by measuring the optical density at 260 and 280 nm. Single-strand cDNA synthesis was carried out with 1 mg of total RNA using SCRIPT Reverse Transcriptase (Jena Bioscience). qPCR analysis of Gli1 was performed with primers 5′ CCAACTCCACAGGCATACAGGAT 3′ and 5′ CACAGATTCAGGCTCACGCTTC 3′ using GoTaq® qPCR Master Mix (Promega).

### Statistical tests

Graphpad Prism 7 was used for applying statistical tests and generating graphs. Two experimental groups were compared using two-sided Student’s t-test while experiments involving more than two experimental groups were analyzed using one- or two-way ANOVA. Number of analyzed cells or experimental replicas for each condition are indicated at figure legend. Error bars indicate standard deviation (SD) and significancy levels were denoted as ns>0.05, * p<0.05, ** p<0.01, *** p<0.001, **** p<0.0001.

## Acknowledgements

We acknowledge the Firat-Karalar lab members for insightful discussions regarding this work. This work was supported by ERC Grant 679140 to ENF, EMBO Installation Grant and Young Investigator Award to ENF, TUBITAK BIDEB 120C148 grant to ENF and Norwegian Cancer Society, Career Development Grant 6839316 to S.P. This project has received funding from the European Union’s Horizon 2020 research and innovation programme under the Marie Sklodowska-Curie grant agreement No 896644 awarded to JD

## Supplementary figures

**Supplementary Figure 1.**
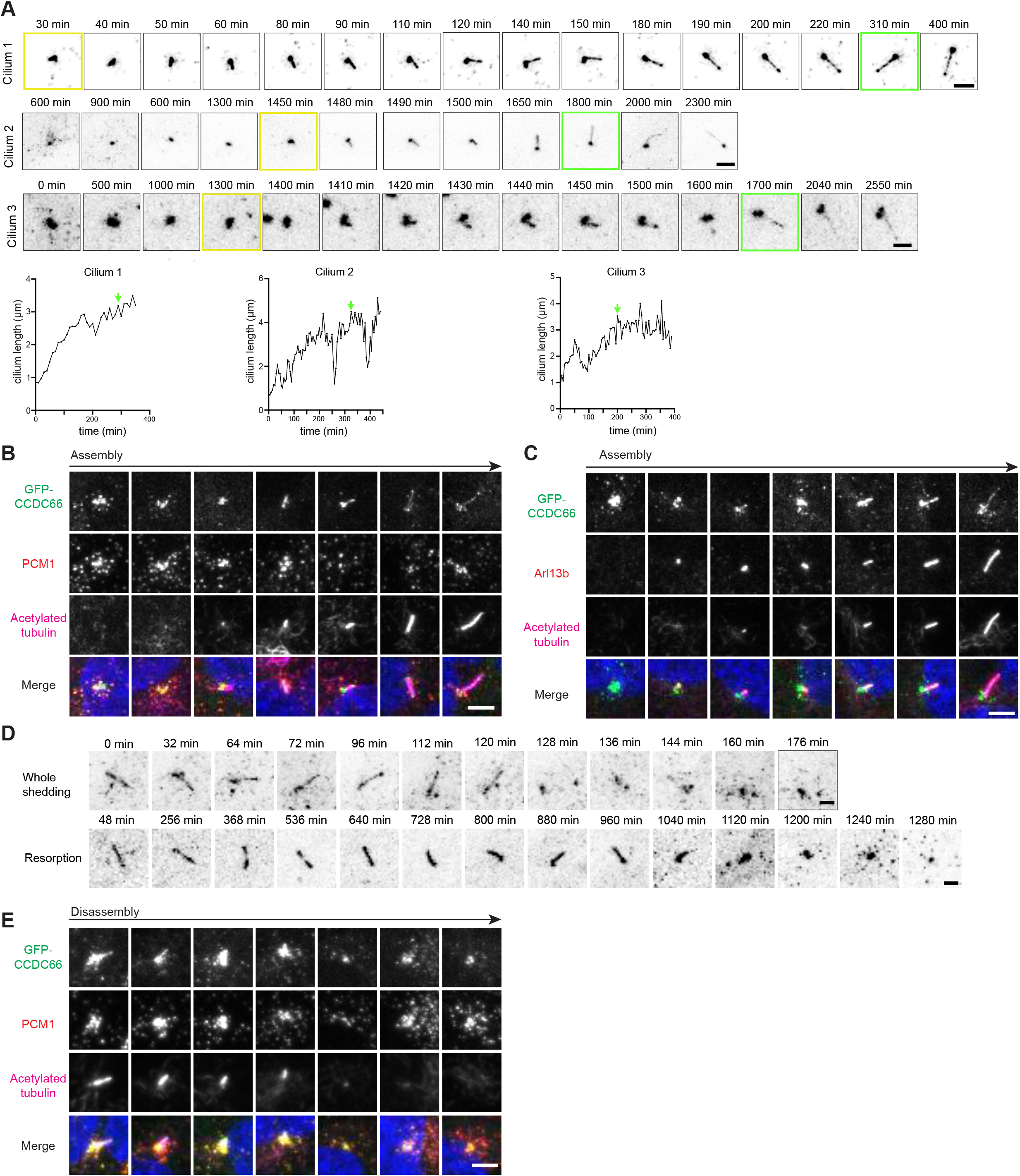
**(A) Spatiotemporal localization dynamics of CCDC66 during cilium assembly**. RPE1::GFP-CCDC66 cells were plated onto Lab-Tek imaging dish at 100% confluency and started to be imaged with confocal microscopy every 10 min immediately after serum starvation. Representative images are from three different cells that form primary cilia. GFP signal is inverted and represented as black onto white background for ciliary and centriolar satellite pools of CCDC66 to be distinguished. Below graphs represent the cilium length over the course of imaging from these three cilia. Cilium length was measured starting from the yellow framed time points. Green framed time points and the green arrow represent reaching the steady state cilium. Scale bar: 2μm **(B and C) CCDC66 localization with respect to centriolar satellite, centrosome and ciliary markers during cilium assembly**. RPE1::GFP-CCDC66 cells were serum starved for 24 hours and fixed with methanol and stained for **(C)** GFP, PCM1, acetylated tubulin or **(D)** GFP, ARL13B and acetylated tubulin antibodies along with DAPI in order to visualize DNA. Images from left to right represent CCDC66 localization during the initiation and elongation phases of primary cilium formation. Scale bar: 5μm **(D) Spatiotemporal localization dynamics of CCDC66 during cilium disassembly**. RPE1::GFP-CCDC66 cells were plated onto Lab-Tek imaging dish at 100% confluency, serum starved for 48 hours to induce cilium formation, and started to be imaged with confocal microscopy every 10 min immediately after serum addition. Representative images are from two cilia that undergo cilium disassembly by whole cilium shedding and resorption. GFP signal is inverted and represented as black onto white background for ciliary and centriolar satellite pools of CCDC66 to be distinguished. Scale bar: 2μm **(E) CCDC66 localization with respect to centriolar satellite, centrosome and ciliary markers during cilium disassembly** After 24h serum starvation, RPE1::GFP-CCDC66 cells were incubated with complete media for 24 hours and fixed with methanol and stained for GFP, PCM1, acetylated tubulin along with DAPI in order to visualize DNA. Images from left to right represent CCDC66 localization during the shortening and disassembly of phases of primary cilium formation. Scale bar: 5μm

**Supplementary Figure 2.**
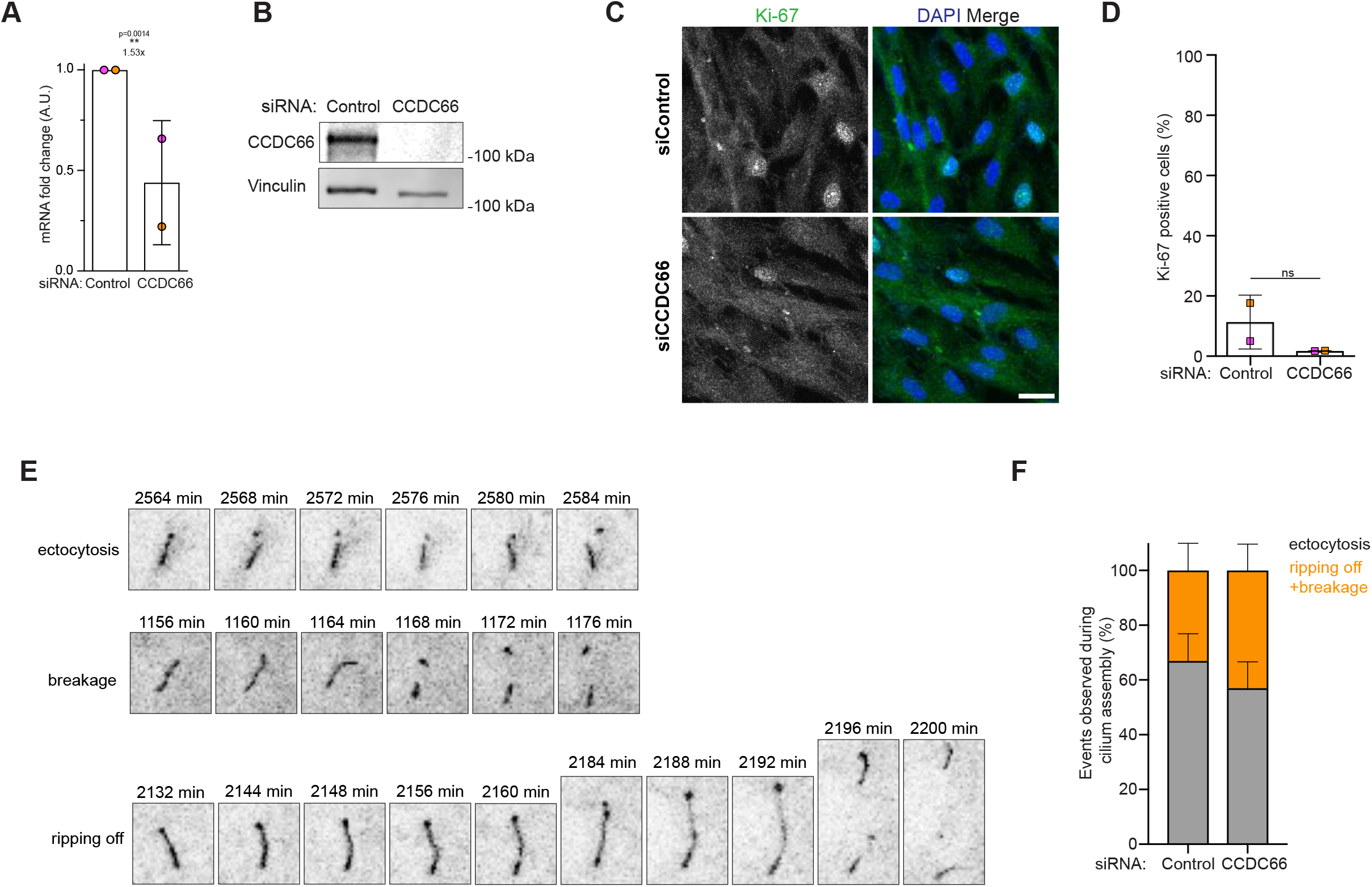
**(A-B) Validation of CCDC66 depletion**. RPE1 cells were transfected with two rounds of control or CCDC66 siRNA. 48 hours later, they were collected and processed for **(A)** qPCR analysis and **(B)** immunoblotting with CCDC66 and vinculin as control loading control. Graph represents CCDC66 mRNA fold change relative to control depletion condition. (*P < 0.5, t-test) **(C and D) Effects of CCDC66 depletion on the percentage of quiescent cells**. RPE1 cells were transfected with two rounds of control or CCDC66 siRNA and serum starved for 24 hours. **(C)** Following fixation with methanol, cells were stained for anti-Ki67 and DAPI for visualization of DNA. **(D)** Graph indicates percentage of Ki-67 positive cells. Data represent the mean ±SD. Scale bar: 15μm. **(E-F) Effects of CCDC66 depletion on steady-state cilia. (C)** Control or CCDC66 depleted RPE1::mCitrine-Smoothened cells were imaged after serum starvation. Cilia from both conditions were observed to undergo ectocytosis, breakage and ripping off events during cilium assembly. Images representing these events are given. **(D)** Graph indicates the percentage of ectocytosis (grey) and breakage+ripping off (orange) events in control and CCDC66 depleted cells. Data represents the mean ±SD. (100 cells/experiment, *P < 0.5, t-test)

**Supplementary Figure 3.**
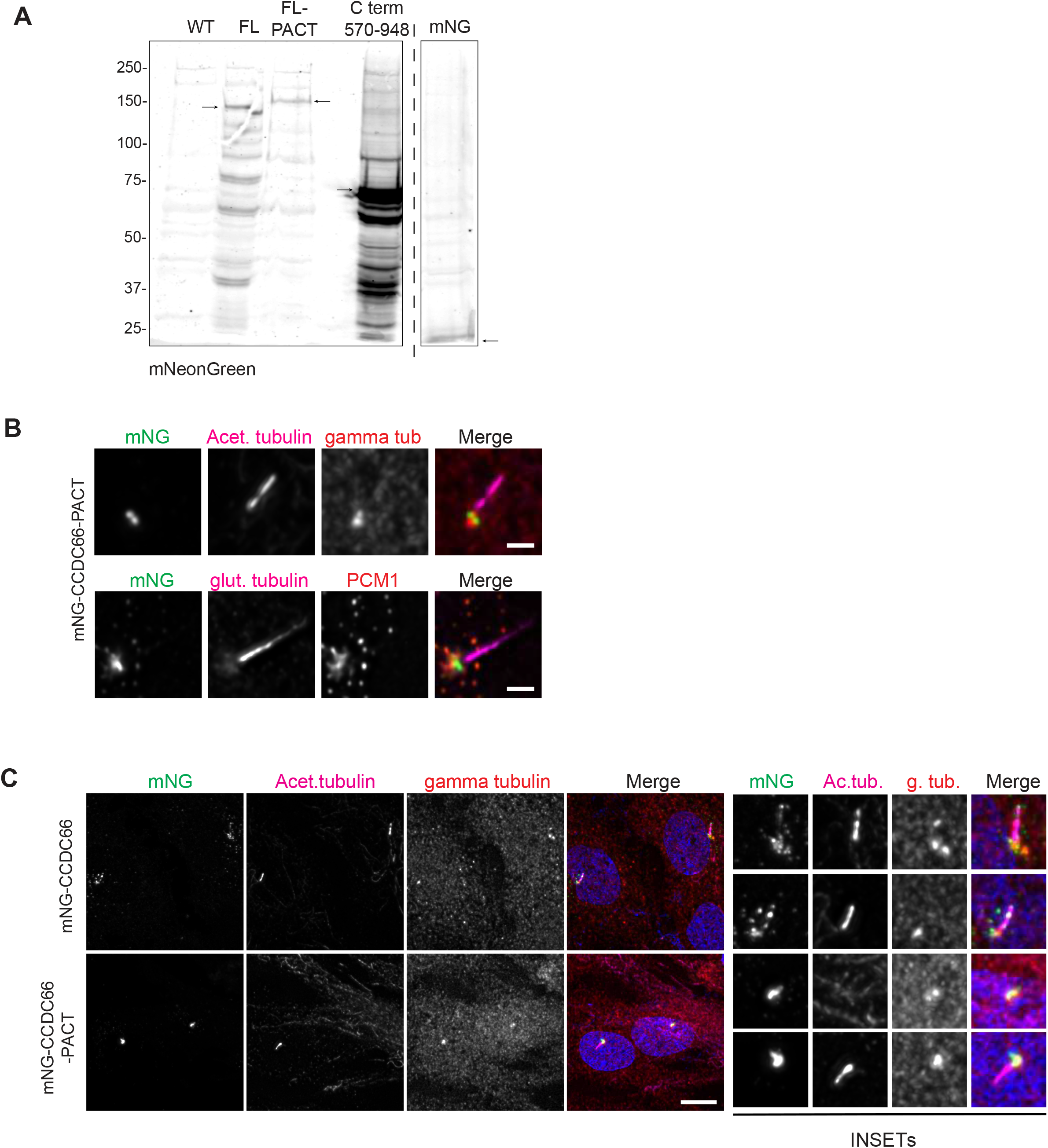
**(A) Validation of stable protein expression**. RPE1 cell lines expressing mNeonGreen, mNeonGreen tagged CCDC66 full length, CCDC66-PACT and CCDC66 C terminal (570-958) were processed and immunoblotted for mNeonGreen. Black arrows indicate the corresponding band for NG tagged protein. **(B-C) mNG-CCDC66-PACT fusion protein localization is restricted to the basal body**. RPE1 cells stably expressing mNeonGreen-CCDC66 full length and mNeonGreen-CCDC66-PACT were serum starved for 48 hours. Following fixation, cells were stained for mNeonGreen, acetylated tubulin, gamma tubulin antibodies and DAPI for visualization of DNA. **(B)** mNG-CCDC66-PACT localization is also represented with images containing single cilium. These cells were also stained for mNeonGreen, polyglutamylated tubulin and PCM1. Insets represent 3x magnification. Scale bar: 2μm for (B) 10 μm for (C).

**Supplementary Figure 4.**
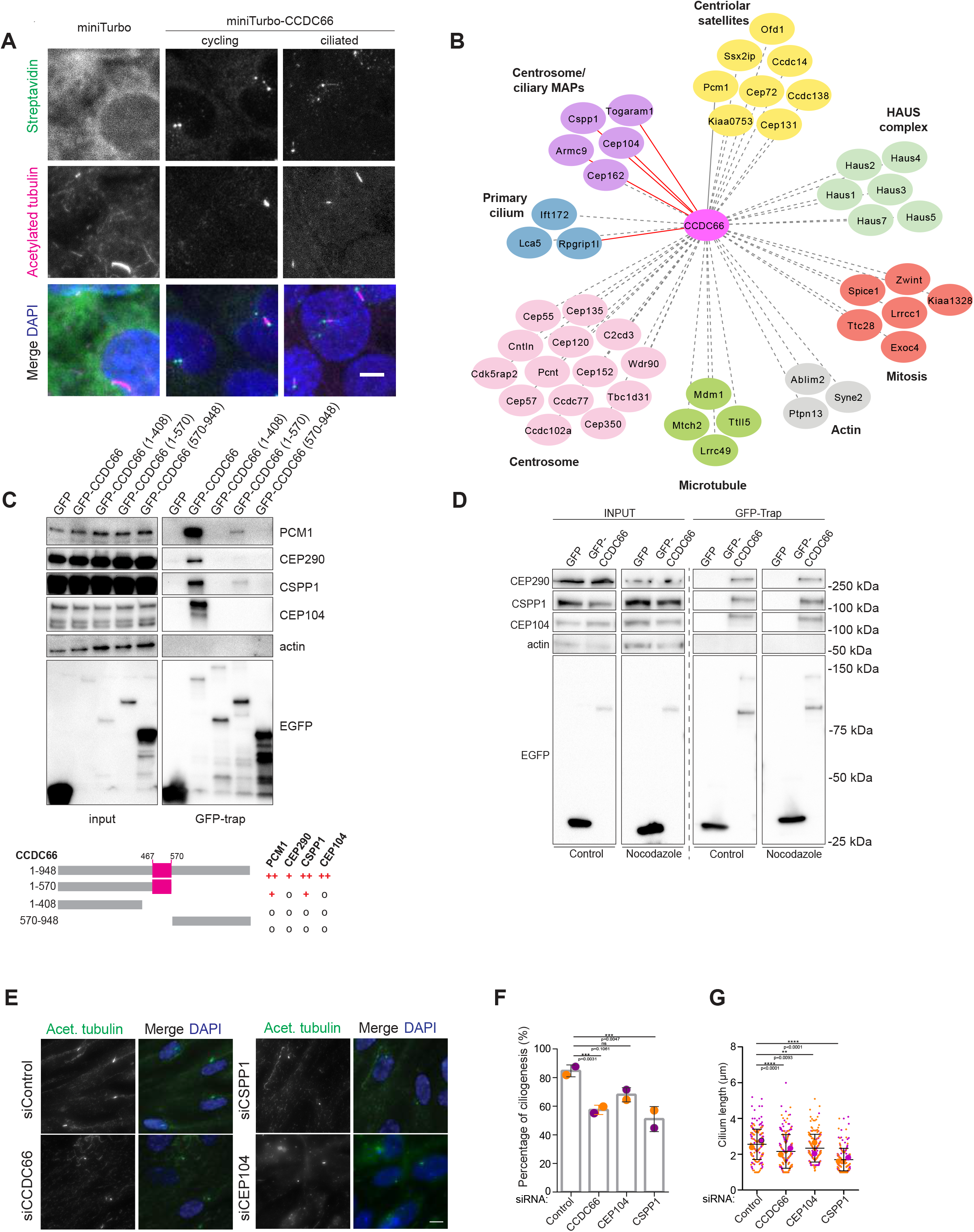
**(A) miniTurbo-CCDC66 induces localized biotinylation at the centrosome, centriolar satellites and primary cilium**. 100% confluency IMCD3 cells stably expressing miniTurboID or miniTurboID-CCDC66 were serum starved for 48 hours. Both cell populations were treated with 500 μM biotin for 30 minutes and fixed with methanol. Cells were stained for anti-acetylated tubulin, Streptavidin and DAPI. Scale bar: 5μm **(B) CCDC66 proximity interaction map in ciliated cells**. Final ciliated CCDC66 proximity interactome list is generated after applying several filtering steps to remove unspecific proteins using NSAF values. Remaining proteins are categorized based on their function and localization information. Dashed line: proximity interaction, solid red line: proximity and direct interactor. **(C) Microtubule-binding fragment of CCDC66 does not interact with CCDC66 full length interactors**. HEK293T cells were transfected with EGFP, EGFP-CCDC66 full length, 1-408, 1-570, 570-948 constructs, and immunoprecipitated with GFP Trap beads. Input and pellet fractions were immunoblotted for GFP, PCM1, CEP290, CSPP1, CEP104 and actin as a loading control. The scheme summarizes CCDC66 N and C terminal constructs and their interactions with indicated proteins (**o**: no interaction, **+**: weak interaction, **++**: moderate interaction, **+++**: strong interaction). **(D) CCDC66 interacts CEP290, CSPP1 and CEP104 independent of microtubules**. HEK293T cells were transfected with EGFP or EGFP-CCDC66 full length. 2 days after transfection, cells were treated with 5 ug/ml nocodazole or 0.02% DMSO as a control. Following incubation, cells were collected, lysed, and subjected to pull down with GFP Trap beads. Input and pellet were immunoblotted with anti GFP, CEP290, CSPP1, CEP104 and actin as a control. **(E-G) Validation of CEP104 and CSPP1 depletion by functional assays**. RPE1 cells were transfected with two rounds of control, CCDC66, CSPP1 or CEP104 siRNAs and serum starved for 48 hours. Following fixation with methanol, cells were stained for acetylated tubulin and DAPI for visualization of DNA. **(F)** Percentage of cilium formation and **(G)** ciliary length were plotted. (100 cell or cilia /experiment, *P < 0.5, **P < 0.01, ***P < 0.001, ****P < 0.0001, ns: not significant, two way ANOVA) Scale bar: 10μm

## Movies

**Movie 1. CCDC66 dynamic localization during cilium assembly** RPE1::mNeonGreen-CCDC66, mScarlet-ARL13B cells were imaged with confocal microscopy every 8 minutes immediately after serum starvation. Scale bar: 5μm

**Movie 2. CCDC66 dynamic localization during cilium disassembly**

After 48 hours serum starvation, RPE1::mNeonGreen-CCDC66, mScarlet-ARL13B cells were imaged with confocal microscopy every 6 minutes immediately upon serum addition. Scale bar: 5-μm

## Table

**Table 1.** Mass spectrometry results of proximity interactors of miniTurboID and miniTurboID-CCDC66 in ciliated IMCD3 cells, related to Supplementary Figure 4A and B. Column explanations of raw data are placed to sheet 2.

## References

Al-Jassar C, Andreeva A, Barnabas DD, McLaughlin SH, Johnson CM, Yu M, van Breugel M. 2017. The Ciliopathy-Associated Cep104 Protein Interacts with Tubulin and Nek1 Kinase. Structure 25: 146–156.

Aydin OZ, Taflan SO, Gurkaslar C, Firat-Karalar EN. 2020. Acute inhibition of centriolar satellite function and positioning reveals their functions at the primary cilium. PLoS Biol 18: e3000679.

Bedoni N, Haer-Wigman L, Vaclavik V, Tran VH, Farinelli P, Balzano S, Royer-Bertrand B, El-Asrag ME, Bonny O, Ikonomidis C et al. 2016. Mutations in the polyglutamylase gene TTLL5, expressed in photoreceptor cells and spermatozoa, are associated with cone-rod degeneration and reduced male fertility. Hum Mol Genet 25: 4546–4555.

Betleja E, Cole DG. 2010. Ciliary trafficking: CEP290 guards a gated community. Curr Biol 20: R928–931.

Blacque OE, Sanders AA. 2014. Compartments within a compartment: what C. elegans can tell us about ciliary subdomain composition, biogenesis, function, and disease. Organogenesis 10: 126–137.

Bodakuntla S, Jijumon AS, Villablanca C, Gonzalez-Billault C, Janke C. 2019. Microtubule-Associated Proteins: Structuring the Cytoskeleton. Trends Cell Biol 29: 804–819.

Braun DA, Hildebrandt F. 2017. Ciliopathies. Cold Spring Harb Perspect Biol 9.

Breslow DK, Holland AJ. 2019. Mechanism and Regulation of Centriole and Cilium Biogenesis. Annu Rev Biochem 88: 691–724.

Broekhuis JR, Leong WY, Jansen G. 2013. Regulation of cilium length and intraflagellar transport. Int Rev Cell Mol Biol 303: 101–138.

Cajanek L, Nigg EA. 2014. Cep164 triggers ciliogenesis by recruiting Tau tubulin kinase 2 to the mother centriole. Proc Natl Acad Sci U S A 111: E2841–2850.

Chien A, Shih SM, Bower R, Tritschler D, Porter ME, Yildiz A. 2017. Dynamics of the IFT machinery at the ciliary tip. Elife 6.

Conkar D, Bayraktar H, Firat-Karalar EN. 2019. Centrosomal and ciliary targeting of CCDC66 requires cooperative action of centriolar satellites, microtubules and molecular motors. Sci Rep 9: 14250.

Conkar D, Culfa E, Odabasi E, Rauniyar N, Yates JR, 3rd, Firat-Karalar EN. 2017. Centriolar satellite protein CCDC66 interacts with CEP290 and functions in cilium formation and trafficking. J Cell Sci.

Conkar D, Firat-Karalar EN. 2020. Microtubule-associated proteins and emerging links to primary cilium structure, assembly, maintenance, and disassembly. FEBS J.

Das A, Dickinson DJ, Wood CC, Goldstein B, Slep KC. 2015. Crescerin uses a TOG domain array to regulate microtubules in the primary cilium. Mol Biol Cell 26: 4248–4264.

Dekomien G, Vollrath C, Petrasch-Parwez E, Boeve MH, Akkad DA, Gerding WM, Epplen JT. 2010. Progressive retinal atrophy in Schapendoes dogs: mutation of the newly identified CCDC66 gene. Neurogenetics 11: 163–174.

den Hollander AI, Koenekoop RK, Mohamed MD, Arts HH, Boldt K, Towns KV, Sedmak T, Beer M, Nagel-Wolfrum K, McKibbin M et al. 2007. Mutations in LCA5, encoding the ciliary protein lebercilin, cause Leber congenital amaurosis. Nat Genet 39: 889–895.

Drivas TG, Bennett J. 2014. CEP290 and the primary cilium. Adv Exp Med Biol 801: 519–525.

Farmer VJ, Zanic M. 2021. TOG-domain proteins. Curr Biol 31: R499–R501.

Firat-Karalar EN, Rauniyar N, Yates JR, 3rd, Stearns T. 2014. Proximity interactions among centrosome components identify regulators of centriole duplication. Curr Biol 24: 664–670.

Frikstad KM, Molinari E, Thoresen M, Ramsbottom SA, Hughes F, Letteboer SJF, Gilani S, Schink KO, Stokke T, Geimer S et al. 2019. A CEP104-CSPP1 Complex Is Required for Formation of Primary Cilia Competent in Hedgehog Signaling. Cell Rep 28: 1907–1922 e1906.

Garcia-Gonzalo FR, Reiter JF. 2017. Open Sesame: How Transition Fibers and the Transition Zone Control Ciliary Composition. Cold Spring Harb Perspect Biol 9.

Gerding WM, Schreiber S, Schulte-Middelmann T, de Castro Marques A, Atorf J, Akkad DA, Dekomien G, Kremers J, Dermietzel R, Gal A et al. 2011. Ccdc66 null mutation causes retinal degeneration and dysfunction. Hum Mol Genet 20: 3620–3631.

Gheiratmand L, Coyaud E, Gupta GD, Laurent EM, Hasegan M, Prosser SL, Goncalves J, Raught B, Pelletier L. 2019. Spatial and proteomic profiling reveals centrosome-independent features of centriolar satellites. EMBO J.

Goncalves J, Pelletier L. 2017. The Ciliary Transition Zone: Finding the Pieces and Assembling the Gate. Mol Cells 40: 243–253.

Graser S, Stierhof YD, Lavoie SB, Gassner OS, Lamla S, Le Clech M, Nigg EA. 2007. Cep164, a novel centriole appendage protein required for primary cilium formation. J Cell Biol 179: 321–330.

Gupta GD, Coyaud E, Goncalves J, Mojarad BA, Liu Y, Wu Q, Gheiratmand L, Comartin D, Tkach JM, Cheung SW et al. 2015. A Dynamic Protein Interaction Landscape of the Human Centrosome-Cilium Interface. Cell 163: 1484–1499.

Gurkaslar HK, Culfa E, Arslanhan MD, Lince-Faria M, Firat-Karalar EN. 2020. CCDC57 Cooperates with Microtubules and Microcephaly Protein CEP63 and Regulates Centriole Duplication and Mitotic Progression. Cell Rep 31: 107630.

He M, Subramanian R, Bangs F, Omelchenko T, Liem KF, Jr., Kapoor TM, Anderson KV. 2014. The kinesin-4 protein Kif7 regulates mammalian Hedgehog signalling by organizing the cilium tip compartment. Nat Cell Biol 16: 663–672.

Jewett CE, Soh AWJ, Lin CH, Lu Q, Lencer E, Westlake CJ, Pearson CG, Prekeris R. 2021. RAB19 Directs Cortical Remodeling and Membrane Growth for Primary Ciliogenesis. Dev Cell 56: 325–340 e328.

Keeling J, Tsiokas L, Maskey D. 2016. Cellular Mechanisms of Ciliary Length Control. Cells 5.

Kobayashi T, Kim S, Lin YC, Inoue T, Dynlacht BD. 2014. The CP110-interacting proteins Talpid3 and Cep290 play overlapping and distinct roles in cilia assembly. J Cell Biol 204: 215–229.

Kohli P, Hohne M, Jungst C, Bertsch S, Ebert LK, Schauss AC, Benzing T, Rinschen MM, Schermer B. 2017. The ciliary membrane-associated proteome reveals actin-binding proteins as key components of cilia. EMBO Rep 18: 1521–1535.

Latour BL, Van De Weghe JC, Rusterholz TD, Letteboer SJ, Gomez A, Shaheen R, Gesemann M, Karamzade A, Asadollahi M, Barroso-Gil M et al. 2020. Dysfunction of the ciliary ARMC9/TOGARAM1 protein module causes Joubert syndrome. J Clin Invest.

Lechtreck KF. 2015. IFT-Cargo Interactions and Protein Transport in Cilia. Trends Biochem Sci 40: 765–778.

Lee J, Chung YD. 2015. Ciliary subcompartments: how are they established and what are their functions? BMB Rep 48: 380–387.

Lu Q, Insinna C, Ott C, Stauffer J, Pintado PA, Rahajeng J, Baxa U, Walia V, Cuenca A, Hwang YS et al. 2015. Early steps in primary cilium assembly require EHD1/EHD3-dependent ciliary vesicle formation. Nat Cell Biol 17: 531.

May EA, Kalocsay M, D’Auriac IG, Schuster PS, Gygi SP, Nachury MV, Mick DU. 2021. Time-resolved proteomics profiling of the ciliary Hedgehog response. J Cell Biol 220.

Mellacheruvu D, Wright Z, Couzens AL, Lambert JP, St-Denis NA, Li T, Miteva YV, Hauri S, Sardiu ME, Low TY et al. 2013. The CRAPome: a contaminant repository for affinity purification-mass spectrometry data. Nat Methods 10: 730–736.

Mick DU, Rodrigues RB, Leib RD, Adams CM, Chien AS, Gygi SP, Nachury MV. 2015. Proteomics of Primary Cilia by Proximity Labeling. Dev Cell 35: 497–512.

Mirvis M, Siemers KA, Nelson WJ, Stearns TP. 2019. Primary cilium loss in mammalian cells occurs predominantly by whole-cilium shedding. PLoS Biol 17: e3000381.

Mirvis M, Stearns T, James Nelson W. 2018. Cilium structure, assembly, and disassembly regulated by the cytoskeleton. Biochem J 475: 2329–2353.

Mukhopadhyay S, Rohatgi R. 2014. G-protein-coupled receptors, Hedgehog signaling and primary cilia. Semin Cell Dev Biol.

Murgiano L, Becker D, Spector C, Carlin K, Santana E, Niggel JK, Jagannathan V, Leeb T, Pearce-Kelling S, Aguirre GD et al. 2020. CCDC66 frameshift variant associated with a new form of early-onset progressive retinal atrophy in Portuguese Water Dogs. Sci Rep 10: 21162.

Nachury MV, Loktev AV, Zhang Q, Westlake CJ, Peranen J, Merdes A, Slusarski DC, Scheller RH, Bazan JF, Sheffield VC et al. 2007. A core complex of BBS proteins cooperates with the GTPase Rab8 to promote ciliary membrane biogenesis. Cell 129: 1201–1213.

Nachury MV, Mick DU. 2019. Establishing and regulating the composition of cilia for signal transduction. Nat Rev Mol Cell Biol.

Nachury MV, Seeley ES, Jin H. 2010. Trafficking to the ciliary membrane: how to get across the periciliary diffusion barrier? Annu Rev Cell Dev Biol 26: 59–87.

Nager AR, Goldstein JS, Herranz-Perez V, Portran D, Ye F, Garcia-Verdugo JM, Nachury MV. 2017. An Actin Network Dispatches Ciliary GPCRs into Extracellular Vesicles to Modulate Signaling. Cell 168: 252–263 e214.

Odabasi E, Batman U, Firat-Karalar EN. 2020. Unraveling the mysteries of centriolar satellites: time to rewrite the textbooks about the centrosome/cilium complex. Mol Biol Cell 31: 866–872.

Odabasi E, Gul S, Kavakli IH, Firat-Karalar EN. 2019. Centriolar satellites are required for efficient ciliogenesis and ciliary content regulation. EMBO Rep.

Patzke S, Redick S, Warsame A, Murga-Zamalloa CA, Khanna H, Doxsey S, Stokke T. 2010. CSPP is a ciliary protein interacting with Nephrocystin 8 and required for cilia formation. Mol Biol Cell 21: 2555–2567.

Pedersen LB, Akhmanova A. 2014. Kif7 keeps cilia tips in shape. Nat Cell Biol 16: 623–625.

Pedersen LB, Schroder JM, Satir P, Christensen ST. 2012. The ciliary cytoskeleton. Compr Physiol 2: 779–803.

Phua SC, Chiba S, Suzuki M, Su E, Roberson EC, Pusapati GV, Schurmans S, Setou M, Rohatgi R, Reiter JF et al. 2017. Dynamic Remodeling of Membrane Composition Drives Cell Cycle through Primary Cilia Excision. Cell 168: 264–279 e215.

Prosser SL, Pelletier L. 2020. Centriolar satellite biogenesis and function in vertebrate cells. J Cell Sci 133.

Quanlong L, Chuanmao Z, Westlake C. 2021. Live-cell fluorescence imaging of ciliary dynamics. Biophysics Reports 9.

Quarantotti V, Chen JX, Tischer J, Gonzalez Tejedo C, Papachristou EK, D’Santos CS, Kilmartin JV, Miller ML, Gergely F. 2019. Centriolar satellites are acentriolar assemblies of centrosomal proteins. EMBO J.

Reiter JF, Leroux MR. 2017. Genes and molecular pathways underpinning ciliopathies. Nat Rev Mol Cell Biol 18: 533–547.

Rezabkova L, Kraatz SH, Akhmanova A, Steinmetz MO, Kammerer RA. 2016. Biophysical and Structural Characterization of the Centriolar Protein Cep104 Interaction Network. J Biol Chem 291: 18496–18504.

Satish Tammana TV, Tammana D, Diener DR, Rosenbaum J. 2013. Centrosomal protein CEP104 (Chlamydomonas FAP256) moves to the ciliary tip during ciliary assembly. J Cell Sci 126: 5018–5029.

Schreiber S, Petrasch-Parwez E, Porrmann-Kelterbaum E, Forster E, Epplen JT, Gerding WM. 2018. Neurodegeneration in the olfactory bulb and olfactory deficits in the Ccdc66 -/- mouse model for retinal degeneration. IBRO Rep 5: 43–53.

Sergouniotis PI, Chakarova C, Murphy C, Becker M, Lenassi E, Arno G, Lek M, MacArthur DG, Consortium UC-E, Bhattacharya SS et al. 2014. Biallelic variants in TTLL5, encoding a tubulin glutamylase, cause retinal dystrophy. Am J Hum Genet 94: 760–769.

Shaner NC, Lambert GG, Chammas A, Ni Y, Cranfill PJ, Baird MA, Sell BR, Allen JR, Day RN, Israelsson M et al. 2013. A bright monomeric green fluorescent protein derived from Branchiostoma lanceolatum. Nat Methods 10: 407–409.

Shannon P, Markiel A, Ozier O, Baliga NS, Wang JT, Ramage D, Amin N, Schwikowski B, Ideker T. 2003. Cytoscape: a software environment for integrated models of biomolecular interaction networks. Genome Res 13: 2498–2504.

Sharp JA, Plant JJ, Ohsumi TK, Borowsky M, Blower MD. 2011. Functional analysis of the microtubule-interacting transcriptome. Mol Biol Cell 22: 4312–4323.

Szymanska K, Johnson CA. 2012. The transition zone: an essential functional compartment of cilia. Cilia 1: 10.

Tanos BE, Yang HJ, Soni R, Wang WJ, Macaluso FP, Asara JM, Tsou MF. 2013. Centriole distal appendages promote membrane docking, leading to cilia initiation. Genes Dev 27: 163–168.

Taschner M, Lorentzen E. 2016. The Intraflagellar Transport Machinery. Cold Spring Harb Perspect Biol 8.

Tu HQ, Qin XH, Liu ZB, Song ZQ, Hu HB, Zhang YC, Chang Y, Wu M, Huang Y, Bai YF et al. 2018. Microtubule asters anchored by FSD1 control axoneme assembly and ciliogenesis. Nat Commun 9: 5277.

Wang J, Barr MM. 2016. Ciliary Extracellular Vesicles: Txt Msg Organelles. Cell Mol Neurobiol 36: 449–457.

Westlake CJ, Baye LM, Nachury MV, Wright KJ, Ervin KE, Phu L, Chalouni C, Beck JS, Kirkpatrick DS, Slusarski DC et al. 2011. Primary cilia membrane assembly is initiated by Rab11 and transport protein particle II (TRAPPII) complex-dependent trafficking of Rabin8 to the centrosome. Proc Natl Acad Sci U S A 108: 2759–2764.

Wheway G, Nazlamova L, Hancock JT. 2018. Signaling through the Primary Cilium. Front Cell Dev Biol 6: 8.

Wiegering A, Ruther U, Gerhardt C. 2018. The ciliary protein Rpgrip1l in development and disease. Dev Biol 442: 60–68.

Wloga D, Joachimiak E, Louka P, Gaertig J. 2017. Posttranslational Modifications of Tubulin and Cilia. Cold Spring Harb Perspect Biol 9.

Yamazoe T, Nagai T, Umeda S, Sugaya Y, Mizuno K. 2020. Roles of TOG and jelly-roll domains of centrosomal protein CEP104 in its functions in cilium elongation and Hedgehog signaling. J Biol Chem 295: 14723–14736.

